# Odd-paired is a late-acting pioneer factor coordinating with Zelda to broadly regulate gene expression in early embryos

**DOI:** 10.1101/853028

**Authors:** Theodora Koromila, Fan Gao, Yasuno Iwasaki, Peng He, Lior Pachter, J. Peter Gergen, Angelike Stathopoulos

## Abstract

Pioneer factors such as Zelda help initiate zygotic transcription in *Drosophila* early embryos, but whether other factors support this dynamic process is unclear. Odd-paired (Opa), a zinc-finger transcription factor expressed at cellularization, controls transition of genes from pair-rule to segmental patterns along the anterior-posterior axis. Finding that Opa also regulates late expression through enhancer *sog_Distal,* along the dorso-ventral axis, we hypothesized that Opa acts as a general timing factor. Chromatin-immunoprecipitation (ChIP-seq) confirmed Opa *in vivo* binding to *sog_Distal* but also identified widespread binding throughout the genome, comparable to Zelda. Furthermore, chromatin assays (ATAC-seq) demonstrate that Opa, like Zelda, influences chromatin accessibility genome-wide, suggesting both are pioneer factors with common as well as distinct targets. Lastly, embryos lacking *opa* exhibit widespread, late patterning defects spanning both axes. Collectively, these data suggest Opa, a general timing factor and likely a late-acting pioneer factor, heralds in a secondary wave of zygotic gene expression.

## INTRODUCTION

The transition from dependence on maternal transcripts deposited into the egg to newly transcribed zygotic transcripts is carefully regulated to ensure proper development of early embryos. During the maternal-to-zygotic transition (MZT), maternal products are cleared and zygotic genome activation occurs (rev. in Hamm and Harrison, 2018; Vastenhouw et al., 2019). In *Drosophila* embryos, the first 13 mitotic divisions involve rapid nuclear cycles (nc), that include only a short DNA replication S phase and no G2 phase, and the nuclei are not enclosed in separate membrane compartments but instead present in a joint cytoplasm (Foe and Alberts, 1983). This streamlined division cycle likely relates to the fast development of *Drosophila* embryos, permitting rapid increase in cell number before gastrulation in a matter of a few hours. Dynamic gene expression is initiated during the early syncytial stage, as early as nc7, and continues to the cellularized blastoderm stage (Ali-Murthy and Kornberg, 2016; Lott et al., 2011). Gene expression patterns may be transient or continuous, lasting through gastrulation or beyond (Kvon et al., 2014). This process is controlled by a specific class of transcriptional factors (TF), called pioneer factors, that bind to closed chromatin cis-regulatory regions to create accessible binding sites for additional transcription factors during development (Iwafuchi-Doi and Zaret, 2014). A pivotal factor in *Drosophila* embryogenesis is the pioneer TF Zelda (Zld), a ubiquitous, maternal transcription factor that binds to promoters of the earliest zygotic genes and primes them for activation (Harrison et al., 2011, 2010; Liang et al., 2008). It was unknown whether a similar regulation exists in other animals, until Pou5f1 (homolog of the mammalian Oct4) was shown to act in an analogous manner to Zld in that it controls zygotic gene activation in vertebrates (Leichsenring et al., 2013).

The exact molecular mechanism that supports widespread activation of zygotic gene expression is not known, but several regulatory mechanisms have been proposed. One model suggests that a decrease in histone levels occurs over time that provides an opportunity for the pioneer factors that herald zygotic gene expression to successfully compete for DNA, in order to access and activate transcription (Hamm and Harrison, 2018; Shindo and Amodeo, 2019). For example, Zld is a pivotal activator of the MZT as it increases accessibility of chromatin at enhancers, to facilitate binding of other transcriptional activators to these DNA sequences and thereby allow initiation of zygotic gene expression (Harrison et al., 2011; Liang et al., 2008; Nien et al., 2011). Zld binds nucleosomes and is considered a pioneer factor (McDaniel et al., 2019). Loss of Zld leads to a large-scale, global decrease in zygotic gene expression as many enhancer regions remain inaccessible and thus non-functional (Schulz et al., 2015). By facilitating chromatin accessibility, Zld has been shown to influence the ability of morphogen transcription factors, Bicoid and Dorsal, to support morphogen gradient patterning (Foo et al., 2014; Xu et al., 2014). While Zld is clearly pivotal for supporting MZT, some genes continue to be expressed even in its absence (Nien et al., 2011). In particular, as chromatin accessibility in the early embryo has recently been shown to be a dynamic process (Blythe and Wieschaus, 2016), it is possible that Zld contributes in a stage-specific manner. It is likely, therefore, that other pioneer factors also exist that contribute to zygotic genome activation.

Furthermore, the embryo undergoes a widespread state change after the 14th nuclear division, and this developmental milestone is classified as the midblastula transition (MBT) (Foe and Alberts, 1983; Shermoen et al., 2010). The cycle slows dramatically and nuclei become cellularized relating to when embryonic programs of morphogenesis and differentiation initiate. Cell membranes emerge to encapsulate nuclei, forming a single layered epithelium. In addition, at nc14, developmental changes relating to DNA replication occur; namely a lengthened S-phase and the introduction of G2 phase. MBT is also associated with clearance of a subset of maternally provided mRNAs, large-scale transcriptional activation of the zygotic genome, and an increase in cell cycle length (Tadros and Lipshitz, 2009; Yuan et al., 2016). We hypothesized that other late-acting pioneer factors manage the MBT in addition or in place of Zld.

An attractive research target in the field of developmental biology is the transcription factor Zinc finger in the cerebellum (ZIC human ortholog); its role in early developmental processes has been established across major animal models and also linked to human developmental pathology (rev. in Aruga and Millen, 2018; Houtmeyers et al., 2013). However, the founding member of the Zic family is the *Drosophila* gene *odd-paired (opa)* (Aruga et al., 1996; Hursh and Stultz, 2018). *opa* is a broadly expressed gene of relatively long transcript length (∼17 kB) that initiates expression at mid-nuclear cycle 14 (Benedyk et al., 1994; Cimbora and Sakonju, 1995) and recently was shown to function as a timing factor in early embryos controlling segmentation (Clark and Akam, 2016). Opa protein has a DNA binding domain containing five Cys2His2-type zinc fingers, and shares homology with mammalian Zic1, 2, and 3 transcription factors. It is categorized as an atypical pair-rule gene, as it is expressed in a broad domain within early embryos rather than the 7-stripe pattern typical of other pair-rule genes. Its pair-rule phenotype relates to its ability to act as a timing factor to control the transition of genes with pair-rule patterns to segmental expression pattern (i.e. from 7-to 14-stripes) (Clark and Akam, 2016). *opa* is a widely expressed gene and has additional functions throughout development. For example, *opa* mutant embryos die before hatching, exhibiting aberrant segmentation, but mutant embryos also exhibit defects in larval midgut formation (Cimbora and Sakonju, 1995). During midgut formation, it regulates expression of a pivotal receptor tyrosine kinase required for proper morphogenesis of the visceral mesoderm (Mendoza-García et al., 2017). In addition, at later stages, Opa supports temporal patterning of intermedial neural progenitors of the *Drosophila* larval brain (Abdusselamoglu et al., 2019). Therefore, *opa* supports a number of patterning roles throughout development, several relating to temporal patterning.

In addition, previous studies had suggested that Opa can influence the activity of other transcription factors to promote gene expression. A well characterized target of Opa in the early embryo is *sloppy-paired 1* (*slp1*), a gene exhibiting a segment polarity expression pattern, for which two distinct enhancers have been identified that are capable of responding to regulation by Opa and other pair-rule transcription factors (Cadigan et al., 1994; Prazak et al., 2010). One of these, the *slp1* DESE enhancer, mediates both Runt-dependent activation and repression (Hang and Gergen, 2017). Opa is clearly a central player in *slp1* regulation by supporting Run’s role as activator at DESE, but its uniform expression pattern does not provide positional information as other pair-rule gene inputs. Furthermore, our recent study showed that Runt regulates the spatiotemporal response of another enhancer, *sog_Distal* (also known as sog_Shadow; Hong et al., 2008; Ozdemir et al., 2011) to support its expression in a broad stripe across the dorsal-ventral (DV) axis on both sides of the embryo (Koromila and Stathopoulos, 2019). Using a combination of fixed and live imaging approaches, our analysis suggested that Run’s role changes from repressor to activator, over time in the context of *sog_Distal*; late expression requires Run input as activator. These analyses of *slp1* DESE and *sog_Distal* regulation supported the view that Opa might provide temporal input into both these enhancers.

The current study was initiated to investigate whether Opa supports late expression through the *sog_Distal* enhancer. Previous studies had not linked Opa to the regulation of DV patterning. Nevertheless, through mutagenesis experiments coupled with live *in vivo* imaging, we provide evidence that Opa does regulate expression of the *sog_Distal* enhancer and that its role is indeed late-acting, occurring in embryos at mid-nc14 onwards whereas the enhancer initiates expression at nc10. Furthermore, using both chromatin immunoprecipitation coupled with high throughput sequencing (ChIP-seq) as well as single-embryo Assay for Transposase-Accessible Chromatin using sequencing (ATAC-seq), our whole-genome data support the view that Opa supports patterning in the embryo by serving as a general timing factor, and possibly a pioneer factor, that broadly influences zygotic transcription at around the time that embryo begin cellularization to support the late phase of the maternal-to-zygotic transition.

## RESULTS

### Opa regulates the sog_Distal enhancer demonstrating a role for this pair-rule gene in DV axis patterning

In a previous study, we created a reporter in which the 650 bp *sog_Distal* enhancer sequence was placed upstream of a heterologous promoter from the *even-skipped* gene (*eve.p*), driving expression of a compound reporter gene containing both tandem array of MS2 sites and the gene *yellow* including its introns (Koromila and Stathopoulos, 2017). The MS2 cassette contains 24 repeats of a DNA sequence encoding an RNA stem-loop when transcribed. The stem-loop is recognized by an MCP-GFP fusion protein, as MCP is a phage protein able to bind the RNA-stem loop whereas GFP provides a strong green signal that can be monitored within the nucleus of *Drosophila* embryos at the site of nascent transcript production associated with the transgene (Bothma et al., 2014). This compound reporter gene allows assay of spatiotemporal gene expression live, through MS2-MCP imaging, or in fixed samples through in situ hybridization to gene *yellow*. We used this *sog_Distal* reporter to assay gene expression in the early embryo, and output was analyzed using a previously defined computational approach to analyze spatiotemporal dynamics (Koromila and Stathopoulos, 2019). Expression through this enhancer initiates at nc10 and continues into gastrulation. In this study, late expression through this enhancer was assayed from nc13 into nc14. Furthermore, nc14 was assayed in four intervals each representing ∼10 min of developmental time: nc14a, nc14b, nc14c, and nc14d, due to its longer length (i.e. ∼45 min compared to ∼15 min for nc13).

In our previous study, mutation of the single Run site in the *sog_Distal* enhancer led to expansion of expression early (i.e. nc13 and nc14a) but loss of the pattern late (i.e. nc14c and nc14d) (Koromila and Stathopoulos, 2019). These results suggested that Run’s role switches from that of repressor to activator in the context of *sog_Distal* enhancer (Koromila and Stathopoulos, 2019). Other studies had suggested that Run can function as either repressor or activator depending on context, as the response of different enhancers to Run is influenced by the presence or absence of other specific transcription factors (Hang and Gergen, 2017; Prazak et al., 2010; Swantek and Gergen, 2004). In particular, Opa is critically required for the Runt-dependent activation of *slp1* (Swantek & Gergen, 2004). We hypothesized that Opa also influences Run’s role in the context of the *sog_Distal* enhancer and thus that Opa functions to support late expression, when Run was identified as providing activating input (Koromila and Stathopoulos, 2019).

Surprisingly, the *sog Distal* 650 bp sequence contains five matches to the 12bp Opa binding site consensus based on vertebrate Zic1/Zic3 when one mismatch is allowed (JASPAR; Figure 1F, G). Upon mutation of these five sites through introduction of 2-4 bp changes to each sequence (i.e. *sogD_ΔOpa*), we found reporter expression as assayed by MS2-MCP live in vivo of nascent transcription was relatively normal up to stage nc14b but, subsequently, exhibited a decrease that is apparent at nc14c (Figure 1B; Movie 1). Using a previously described quantitative analysis pipeline (Koromila and Stathopoulos, 2019), we quantified the MS2-MCP signal in embryos containing either the wildtype *sog_Distal* or *sogD_ΔOpa* reporters, confirming that Opa expression is greatly reduced at nc14c for the mutant reporter compared to wildtype (Figure 1 - supplement figure 1E). A similar loss of late expression only (i.e. nc14c onwards) was obtained when even a single Opa site is mutated (Figure 1 - figure supplement 1A-C) comparable to when the Run site is mutated (Koromila and Stathopoulos, 2019). These results support the view that Opa promotes expression through *sog_Distal* from nc14c onwards at least in part by serving to switch Run from repressor to activator (see Discussion).

**Figure 1.**
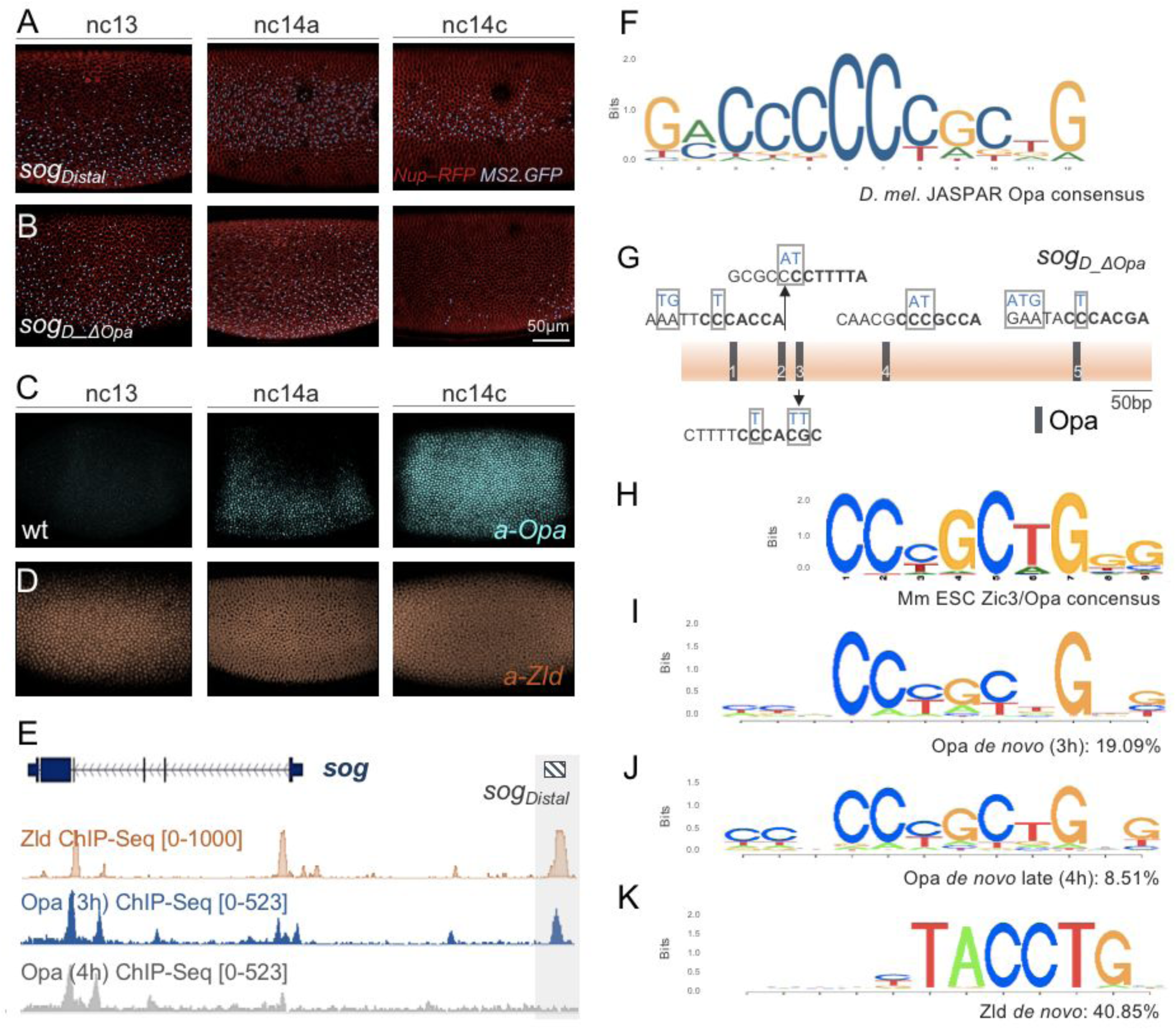
Opa is required to support activation of reporter expression at late nc14, just preceding gastrulation. In this and all other subsequent figures lateral views of embryos are shown with anterior to the left and dorsal side up, unless otherwise noted. (A, B) Stills from representative movies of the two indicated *sog_Distal* MS2-yellow reporter variants (A) *sog_Distal* (A) or *sogD_DOpa* (B) in which 5 predicted Opa binding sites were mutated as shown in (G) to detect transcription via MS2-MCP-GFP (Koromila and Stathopoulos, 2019) imaging at three representative time points: nc13, nc14a, and nc14c. Blue dots for each construct indicate presence of GFP+ dots, representing nascent transcripts labeled by the MS2-MCP system; thresholding was applied and remaining signals identified by the Imaris Bitplane software, for visualization purposes only. Nuclei were labelled by Nup-RFP (Lucas et al., 2018). Scale bar represents 50μm. (C, D) Anti-Opa (C) and anti-Zld (D) antibody staining of early wild-type embryos at nc13, nc14a and nc14c. (E) Integrative Genomics Viewer (IGV) genome browser track of the *sog* locus showing Zld ChIP-seq data for embryos of stage 5 (GEO dataset GSM763061: nc13-nc14) in comparison with Opa ChIP-seq for embryos of two stages: 3h (stage 4/5) and 4h (stage 6/7) (see Methods). Zld and Opa 3h ChIP-seq samples are of overlapping timepoints, whereas Opa 4h ChIP-seq sample is later. Peaks of predicted binding are detected at the *sog_Distal* enhancer and weak binding is detected at the *sog_Intronic* enhancer (Markstein et al., 2002) and another putative enhancer in the third intron. Gray box marks the *sog_Distal* enhancer location that is, bound by both Zld and Opa (3h) early, but not bound late (Opa 4h). (F) JASPAR consensus binding site for Opa based on mammalian Zic proteins identified using SELEX (Khan et al., 2018; Sen et al., 2010). (G) Five matches to the Jaspar consensus binding site for Opa (F) are present within the 650bp *sog_Distal* enhancer region; 12 bp matches shown (black) allowing for 1 mismatch to consensus. These predicted Opa sites were mutated by introduction of changes (blue bases above sequence) predicted to eliminate binding creating *sogD_Δopa* (B; see Methods). Bases in bold (7bp) indicate matches to the Opa *de novo* motifs identified by ChIP-seq analysis (see I,J). For sake of comparison to consensus sequence, a subset of sites are shown as reverse complement. (H) Consensus binding site for mus musculus Zic3/Opa homolog identified using ChIP-seq (Lim et al., 2010). (I-K) Sequence logo representations of consensus binding site identified as the most significant site in the Opa 3 h(I) 3h, Opa 4h (J), or Zld (K) ChIP-seq datasets defined by HOMER *de novo* motif analysis (Central motif enrichment P-values 1e-566, 1e-354, and 1e-3283, respectively).

The timing of Opa expression supports a role for this factor in driving expression of *sog_Distal* at mid-nc14, approximately at the time of the MBT. Using an anti-Opa antibody (Mendoza-García et al., 2017), we examined spatiotemporal dynamics associated with Opa protein in the early embryo through analysis of localization in a time series of fixed embryos. Opa expression is absent at nc13, first observed at nc14a, and encompasses its mature full pattern approximately half-way into nc14, at nc14c (Figure 1C). The timing of Opa onset of expression correlates with the timing of loss of late expression from the *sog_Distal* reporter, observed when Opa sites are mutated (Figure 1C, compare with 1B). On the other hand, the ubiquitous, maternal transcription factor Zld is detected throughout this time period including at nc13 (Figure 1D). Loss of Zld input to *sog_Distal* through mutagenesis of Zld sites, contrastingly, leads to decreased spatial extent of the reporter pattern (sog_Shadow; Yamada et al., 2019) rather than an overall loss of expression as observed when Opa binding sites are mutated (Figure 1B).

These results suggested that Opa may function as a timing factor, to regulate expression of *sog_Distal*, specifically, mid nc14 and later. Recent studies have demonstrated that the *opa* gene is generally important for the temporal regulation of segmental patterning in *Drosophila* as well as in *Tribolium* (Clark and Akam, 2016; Clark and Peel, 2018). However, as our results suggest a role for Opa in the regulation of *sog* expression, which relates to DV patterning, we hypothesized Opa’s role extends beyond control of segmentation along the AP axis to patterning of the embryo, in general.

### Use of anti-Opa antibody to conduct assay of in vivo occupancy to DNA through ChIP-seq analysis

To examine the *in vivo* occupancy of Opa in early *Drosophila* embryos, we conducted chromatin immunoprecipitation coupled to high throughput sequencing (ChIP-seq). Anti-Opa antibody was used to immunoprecipitate chromatin obtained from two embryo samples of average age 3h (roughly stages 4-6) or 4h (roughly stages 5-7) (see Methods). Focusing on the Opa 3h ChIP-seq dataset, the MACS2 peak caller was used to identify 16,085 peaks, representing an estimate of the number of genomic positions that are occupied by Opa *in vivo*. 200 bp-regions associated with these peaks were analyzed using the HOMER program (Heinz et al., 2010) to identify overrepresented sequences that relate to transcription factor binding sites (see Methods). The most significant hit, present in over 19% of all peaks, is a 7 bp core sequence exhibiting partial overlap with the 12 bp Opa JASPAR site (Figure 1I, compare with 1F) as well as sharing significant homology with the binding site observed for mammalian homolog Zic transcription factors (e.g. see Figure 1H; Lim et al., 2010); however, this site exhibits less overlap with consensus binding site defined for Opa *in vitro* by systematic evolution of ligands by exponential enrichment (SELEX) (Sen et al., 2010). A second motif exhibiting extended homology with the JASPAR Opa consensus was also identified through analysis of the 3h ChIP-seq dataset, but this extended site is present at lower abundance (Figure 2 - supplemental figure 1B). In the *sog_Distal* enhancer sequences, the five initial matches made to the JASPAR Opa binding site consensus also match the Opa *de novo* ChIP-seq derived consensus in 6 of the 7 bases. However, there is a notable mismatch in the 3’ most position; while the *de novo* Opa consensus at 3h does not include A in this position, both the JASPAR site and *de novo* Opa consensus derived for the 4h ChIP-seq timepoint do (Figure 1F, J, compare with 1G). It is possible that the predominant Opa *de novo* motifs derived from the ChIp-seq datasets represent high affinity sites, whereas the five Opa sites found in *sog_Distal* represent lower-affinity, cooperating sites; alternatively, the *de novo* Opa consensus derived from ChIP-seq may relate to a particular heterodimeric complex that is present only *in vivo* in the embryo at the assayed stages.

We hypothesized that Opa might also support expression of *sog_Distal* late, specifically, following mid-nc14. Independent chromatin immunoprecipitation experiments have analyzed Zld *in vivo* binding at a similar stage (i.e. nc13-nc14 early is roughly equivalent to 3h) and detected widespread binding of Zld throughout the genome including at the *sog_Distal* sequence (Figure 1E) (Foo et al., 2014; Harrison et al., 2011). We compared Opa binding to *sog_Distal* as well as to other regions through comparison of Opa 3h or 4h ChIP-seq experiments. Opa ChIP-seq detects Opa occupancy at the *sog* locus, including at the *sog_Distal* enhancer sequence, in the 3h but not the 4h time point (Figure 1E). At gastrulation, *sog_Distal* enhancer output changes from a broad lateral stripe to a thin stripe expressed at the midline; it is possible that at this stage, expression of *sog_Distal* is no longer directly dependent on Opa.

### Assay of overrepresented sites associated with Opa ChIP-seq peaks

Region overlap analysis was used to identify overlapping regions of occupancy for Opa (3h) and Zld using the respective ChIP-seq datasets obtained independently but assaying embryos of similar stage (see Methods). Each ChIP-seq experiment identified 16,085 peaks of occupancy representing locations in the genome that are occupied by either Opa or Zld. Opa and Zld have 6087 peaks in common (Opa-Zld overlap), whereas 9998 regions were bound by Opa alone (Opa-only) and 10781 regions were bound by Zld alone (Zld-only) (Figure 2D). When the Opa and Zld binding site consensus sequences, generated *de novo* by assay of the full ChIP-seq datasets (i.e. Figure 1I,K), are used to analyze binding in each of these classes, it is clear that these motifs are differentially enriched in the Opa-only (Figure 2A), Zld-only (Figure 2B), and Opa-Zld overlap regions (Figure 2C) (also see Figure 2 - supplemental figure 1).

**Figure 2.**
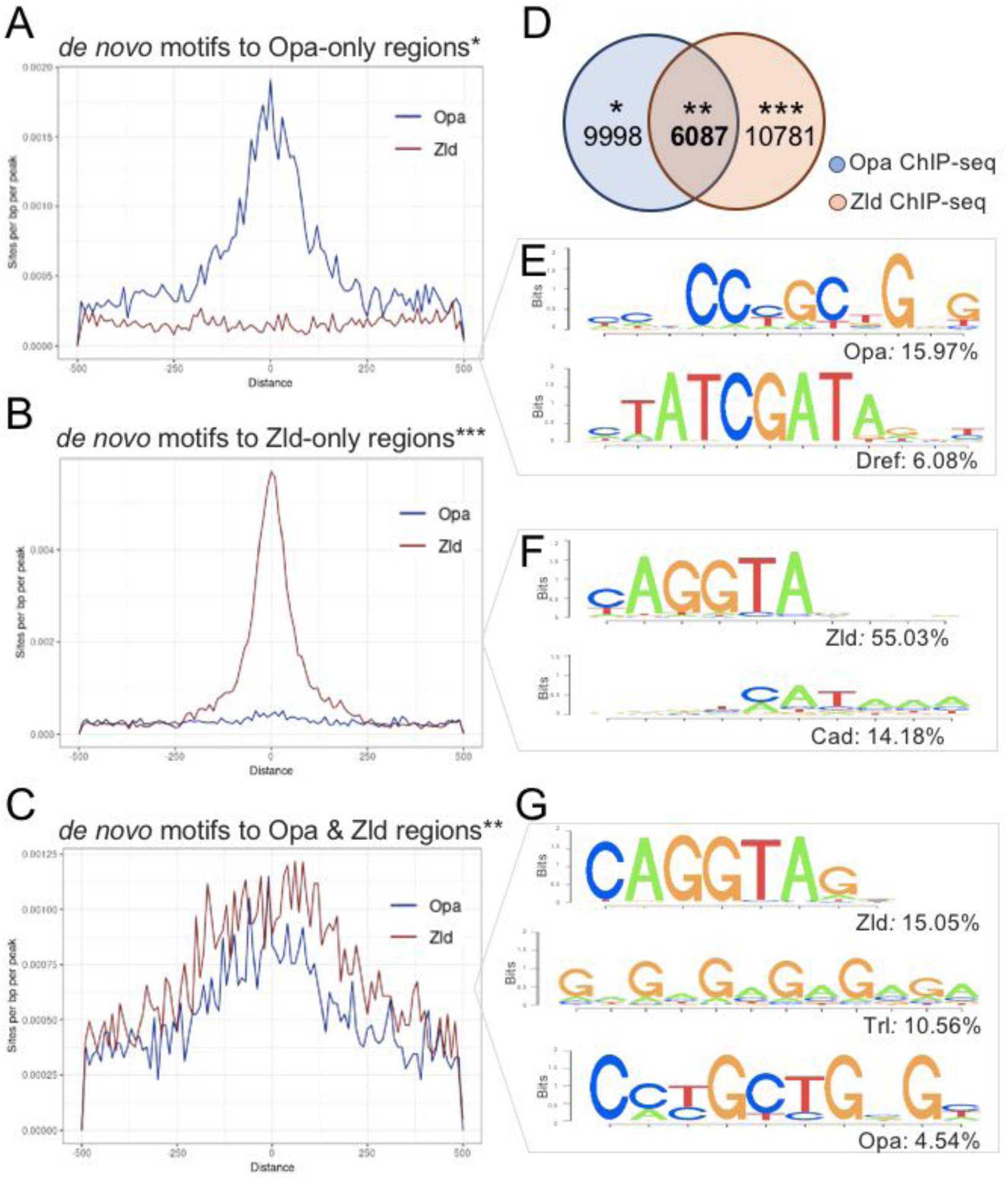
ChIP-Seq analysis of Opa chromatin occupancy in the early *Drosophila* embryo (nc14 to gastrulation). (A-C) Aggregation plots showing enrichment of top *de novo* motifs identified from Opa (3h) peaks and Zld peaks surrounding. (A) Opa_only bound regions (after exclusion of Zld-only and Opa-Zld overlap peaks); (B) Zld-only bound regions (after exclusion of Opa and Opa-Zld overlap peaks); and (C) for Opa-Zld overlap regions. (D) Venn diagram of ChIP-seq data from *Drosophila* embryos during cellularization showing the number of peaks called using MACS2 through analysis of: Opa ChIP-seq (blue) and Zld ChIP-seq (orange) data. Opa and Zld experiments used embryos of 2.5-3.5h in age (stage 5) or nc13/nc14 early, which are overlapping timepoints. Opa-only peaks (*); Opa-Zld overlap peaks (**); Zld-only peaks (***). (E-G) Sequence logo representations of 2-3 most abundant motifs identified using HOMER *de novo* motif analysis within the three corresponding sets of peaks (A: Opa-only, B: Zld-only, C: Opa-Zld overlap peaks). As expected, motifs exhibiting match to the JASPAR Opa and Zld consensus sequences are found. In addition, motif exhibiting match to Dref (E), Caudal (Cad; F), and Trl/GAF (G) were also uncovered in particular datasets. Sequence logo height indicates nucleotide frequency; corresponding Percentage of peaks containing match to motifs also shown for each set, as indicated. P-values representing the significance of motifs’ enrichment compared with the genomic background are as follows: 1) Opa-only peaks: Opa motif enrichment P-value=1e-522; Dref motif enrichment P-value=1e-119, 2) Zld-only peaks: Zld motif enrichment P-value=1e-2833, Caudal motif enrichment P-value=1e-63, 3) Opa-Zld overlap peaks: Zld motif enrichment P-value=1e-240; insulator Trl/GAF motif enrichment P-value=1e-76; Opa motif enrichment P-value=1e-43 (see also Figure 2 - supplemental table 1).

The HOMER sequence analysis program was used to identify overrepresented motifs within these three classes of peaks: Opa-only, Zld-only, or Opa-Zld overlap; in order to identify associated motifs that might provide insight into the differential or combined functions of Opa and Zld. As expected, the top motifs in each class matched the Opa or Zld consensus sequences with 16% of the Opa-only peaks containing an Opa site; 55% of the Zld-only peaks containing a Zld site; and 5% and 15% of the Opa-Zld overlap peaks containing an Opa or Zld site, respectively (Figure 2E-G). In addition, the second-most significant site identified in each class of called peaks corresponds to Dref (6%) for Opa-only; Caudal (Cad; 14%) for Zld-only; and Trl/GAF (11%) for Opa-Zld overlap (Figure 2E-G). Dref (DNA replication-related element-binding factor) is a BED finger-type transcription factor shown to bind to the sequence 5′-TATCGATA (Hirose et al., 1993), a highly conserved sequence in the core promoters of many *Drosophila* genes (Ohler et al., 2002); whereas Cad encodes a homeobox transcription factor that is maternally provided but forms a concentration-gradient enriched at the posterior (Mlodzik et al., 1985) and exhibits preferential activation of DPE-containing promoters (Shir-Shapira et al., 2015). These sites and other additional, significant co-associated sites were uncovered when all called peaks within samples [i.e. Opa (3h), Opa (4h), and Zld] were analyzed by HOMER including a highly significant unknown motif associated with both Opa samples (Figure 1 - figure supplement 2).

Collectively, these results support the view that Zld acts early whereas Opa acts later, and that distinct sets of transcription factors likely help each of these factors to support their different roles (see Discussion).

### Opa ChIP-seq peaks are associated with late-acting enhancers driving expression along both axes

A direct comparison of the Opa (3h) and Zld ChIP-seq occupancy at stage 5, through aggregation plots, suggests that the two transcription factors can bind either simultaneously to the same enhancers (e.g. Figure 2C) or independently to distinct enhancers (e.g. Figure 2A,B). Furthermore, the respective sites appear explanatory for the observed *in vivo* occupancy to DNA sequences as the matches to the consensus sequences correlate with the center of the peak (Figure 2A-C). Furthermore, the widespread binding of Opa in the genome supports the view that this factor functions broadly to support gene expression. Within the total set of 16085 Opa chromatin immunoprecipitation peaks associated with the 3h timepoint, we found, surprisingly, that Opa is associated with genes expressed along both axes: anterior-posterior (AP; Figure 3A-D) and dorsal-ventral (DV; Figure 3E-G), including but not limited to genes involved in segmentation (e.g. *slp1* and *oc*: Figure 3C,D) as predicted by previous studies (e.g. Clark and Akam, 2016; Prazak et al., 2010). We also found evidence in occupancy trends for Opa and Zld that suggest these factors both influence the timing of enhancer action.

**Figure 3.**
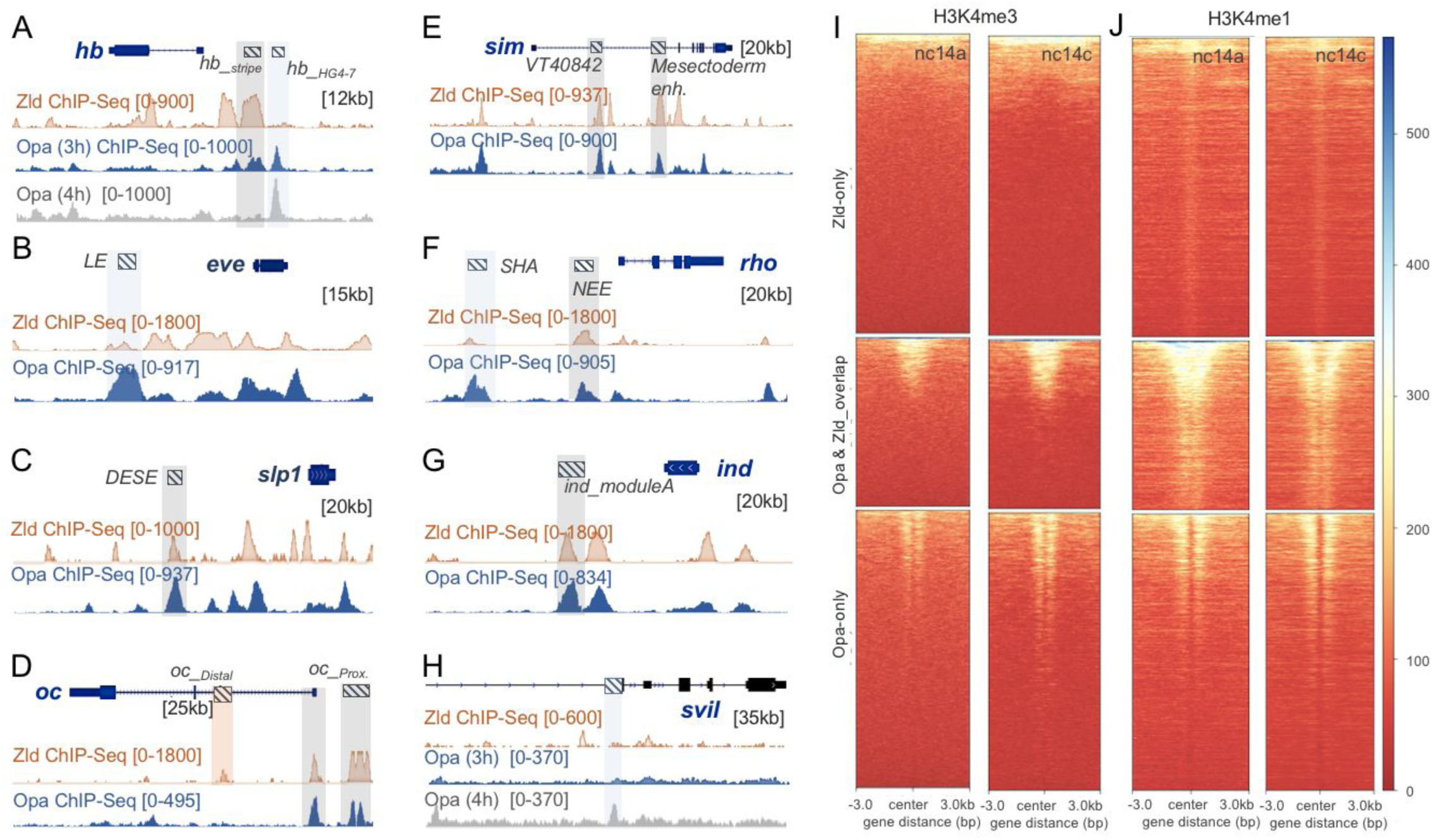
Opa chromatin immunoprecipitation (ChIP) demonstrates global binding at the early *Drosophila* embryo during cellularization. (A-H) IGV browser tracks of genes expressed along either the AP (A-D) or DV (E-H) axes showing single Opa (blue) and Zld (orange) ChIP-seq replicates (as indicated). Anti-Opa antibody was used to immunoprecipitate chromatin isolated from embryos ∼3 hours in age (see Methods). Published Zld ChIP-seq data for stage 5 embryos is shown as a point of comparison (GSM763061: nc13-nc14; Harrison et al., 2011). Gray boxes indicate regions with significant occupancy by both Opa and Zld as detected by ChIP peaks, which can be located at either promoter and/or distal regions (A, C-E, G, H). Light blue boxes indicate regions with significant Opa-only binding at promoter and/or distal regions (B, F). (I, J) Heatmaps produced by deepTools (see Methods) were used to plot histone H3K4me3 (I) and H3K4me1 (J) signal intensities around different ChIP-seq regions (Zld-only, Opa-Zld overlap, and Opa-only bound ChIP-Seq peaks) at two different timepoints nc14a and nc14c (stage 5). Heatmaps are centered on ChIP-seq peak summits. Key indicates histone signal intensities (deepTools normalized RPKM with bin size 10) around different ChIP-seq regions.

Enhancers enriched for Opa-occupancy tend to be active starting at nc14, at the earliest, whereas earlier acting enhancers tend to be bound by Zld alone or Zld and Opa together (Figure 3A-G). Opa binding (3h) is associated with enhancers that are initiated in nc14: *hb_stripe* (Figure 3A; Koromila and Stathopoulos, 2017; Perry et al., 2012), *slp1_DESE* (Figure 3C; Prazak et al., 2010), *oc_Prox* (Figure 3D; Chen et al., 2012; Perry et al., 2011), and *ind_moduleA* (Figure 3G; Stathopoulos and Levine, 2005a). Additionally, Opa binding is also associated with enhancers that are activated later at gastrulation: *hb_HG4-7* (Figure 3A; Hirono et al., 2012), *eve_LE* (Figure 3B; Fujioka et al., 1996), *sim_VT40842* (Figure 3E; Kvon et al., 2014), and *rho_SHA* (Figure 3F; Pearson and Crews, 2014; Rogers et al., 2017). In particular, the *hb_HG4-7* enhancer is not bound by Zld early but is bound by Opa increasingly from 3h to 4h timepoint (Figure 3A); in contrast, the *hb_stripe* enhancer, which is active earlier at nc14, is strongly bound by Zld and also exhibits a concomitant reduction in occupancy of Opa from the 3h to 4 h timepoint. Similarly, the *eve_*LE enhancer is bound by Opa, with little input from Zld, and is also late-acting relative to all other *even-skipped* (*eve*) stripe enhancers, as it mediates the refinement of all seven *eve* stripes after they are initially positioned (Figure 3B; Fujioka et al., 1996). The *SHA* enhancer associated with the gene *rhomboid* (*rho_SHA*) is bound by Opa predominantly and drives expression in the midline at gastrulation (Figure 3F); in contrast, the *rho_NEE* enhancer is bound by both Opa as well as Zld supports *rho* expression earlier in the cellularized blastoderm (Ip et al., 1992). Collectively, these results show that Opa binds to enhancers known to function at nc14 (stage 5) or later, at or following gastrulation (e.g. stage 6/7), and, in contrast, enhancers that receive input from Zld-alone or Opa+Zld tend to be active earlier. However, we can not dismiss a role for Zld at later stages, as the VT40842 enhancer associated with the gene *single-minded* (*sim*) is active later (stage 6) but is bound by both Opa and Zld at an earlier stage (Figure 3E).

To assay whether Opa binding changes in time, we compared Opa occupancy in the two ChIP-seq samples representing two timepoints, 3h (Opa_early) and 4h (Opa_late). There are 9995 peaks in common, 5859 peaks specifically associated with Opa_early, and 6657 peaks specifically associated with Opa_late suggesting changing targets over time (Figure 3 - supplemental figure 1C,D). For example, *sog_Distal* and *hb_stripe* enhancers are Opa-target only bound at the early (3h) timepoint (Figure 1E and 3A); whereas, in turn, *Supervillin* (*Svil*) is an Opa-target bound predominantly at the late (4h) timepoint (Figure 3H). However, the position of peaks, generally, does not appear to change much over time, as most Opa peaks continue to be localized in the same positions at both timepoints (Figure 3 - supplemental figure 1B).

Furthermore, Opa associates with promoters whether or not Zld is co-associated. While Zld was shown to preferentially associate to promoters at earlier time points (nc8; Harrison et al., 2011), we found that binding of Zld to Zld-only enhancers occurs in more distal regions at stage 5 (Figure 3 - supplemental figure 1A). It is possible that once Opa is expressed it preferentially associates with promoter regions and either competes and/or co-regulates with Zld (see Discussion).

### H3K4me3 and H3K4me1 histone marks at nc14 are enriched at regions occupied by Opa

Previous genomics studies have demonstrated that particular histone marks correlate with active enhancers at different developmental stages in the early embryo. For example, there is a dramatic increase in the abundance of histone modifications at the MZT, coinciding with zygotic genome activation (Schulz and Harrison, 2019). We investigated whether Opa-only, Zld-only, and Opa-Zld overlap regions (Figure 2D) exhibit differences in chromatin marks that might support our working hypothesis that Opa-associated regions are active later than Zld-only regions. For the purposes of this analysis, ten published ChIP-seq datasets relating to histone/histone modifications were investigated to assay coincidence of any marks with Opa and/or Zld-bound regions identified by our analysis (see Methods). Only H3K4me3 and H3K4me1 were found to exhibit a difference in Opa versus Zld bound peaks (Figure 3I,J). Both histone marks are first detectable at the MBT, while absent prior to nc14a; whereas their associated genes are considered to be activated at later stages, for example, at MBT following early zygotic activation (Chen et al., 2013; Li et al., 2014).

Heatmap modules of deepTools (see Methods) were used to calculate and plot histone H3K4me3 and H3K4me1 signal intensities assayed at two timepoints, nc14a and nc14c, at different sets of ChIP-seq defined regions: Opa-only; Zld-only; or Opa-Zld overlap. Our analysis shows that Zld-bound regions are underrepresented for H3K4me3, as shown previously (Li et al., 2014), as well as exhibit less enrichment of H3K4me1 at both time points relative to Opa or Opa-Zld overlap bound regions (Figure 3I,J). To our surprise, Opa-Zld overlap and Opa-only bound regions are more enriched for both H3K4me3 and H3K4me1, compared to the Zld-only bound regions (Figure 3I,J). The unique histone methylation pattern exhibited by these classes of peaks, in particular, indicates that Opa targets are likely subject to later developmental regulation. The higher levels of H3K4me1 in the Opa-bound peaks could potentially also be connected to a poised state of late-acting enhancers (Koenecke et al., 2017) or because these regions correspond to other cis-regulatory elements, promoters or insulators, for example (Bonn et al., 2012; Rada-Iglesias et al., 2011) (see Discussion).

### Global changes in chromatin accessibility result upon knock-down of Opa

We hypothesized that Opa functions as a pioneer factor to regulate temporal gene expression starting at nc14 in the celluarizing blastoderm. To test this view, we investigated whether Opa functions to regulate chromatin accessibility. We used ATAC-seq (Assay for Transposase-Accessible Chromatin using sequencing) to investigate the state of chromatin accessibility in embryos that had been depleted of *opa* in comparison to wild type. To start, embryos were depleted of Opa by expression of a short hairpin (sh) RNAi construct (Staller et al., 2013) that we show is effective in reducing *opa* transcript levels when expressed at high levels maternally and zygotically in embryos (Figure 4A,B). In a side by side comparison, we found that *opa* RNAi embryos exhibit phenotypes similar to *opa^1^* zygotic mutants (Figure 4 - supplemental figure 1B, C). However, *opa* expression is retained, though slightly reduced, in embryos in which *zld* levels are knocked-down, using a similar approach, through *zld* RNAi (i.e. *sh zld*; Figure 4C). This results suggests that *opa* expression is not completely under Zld regulation, and indicates these factors could have separable roles (see Discussion).

**Figure 4.**
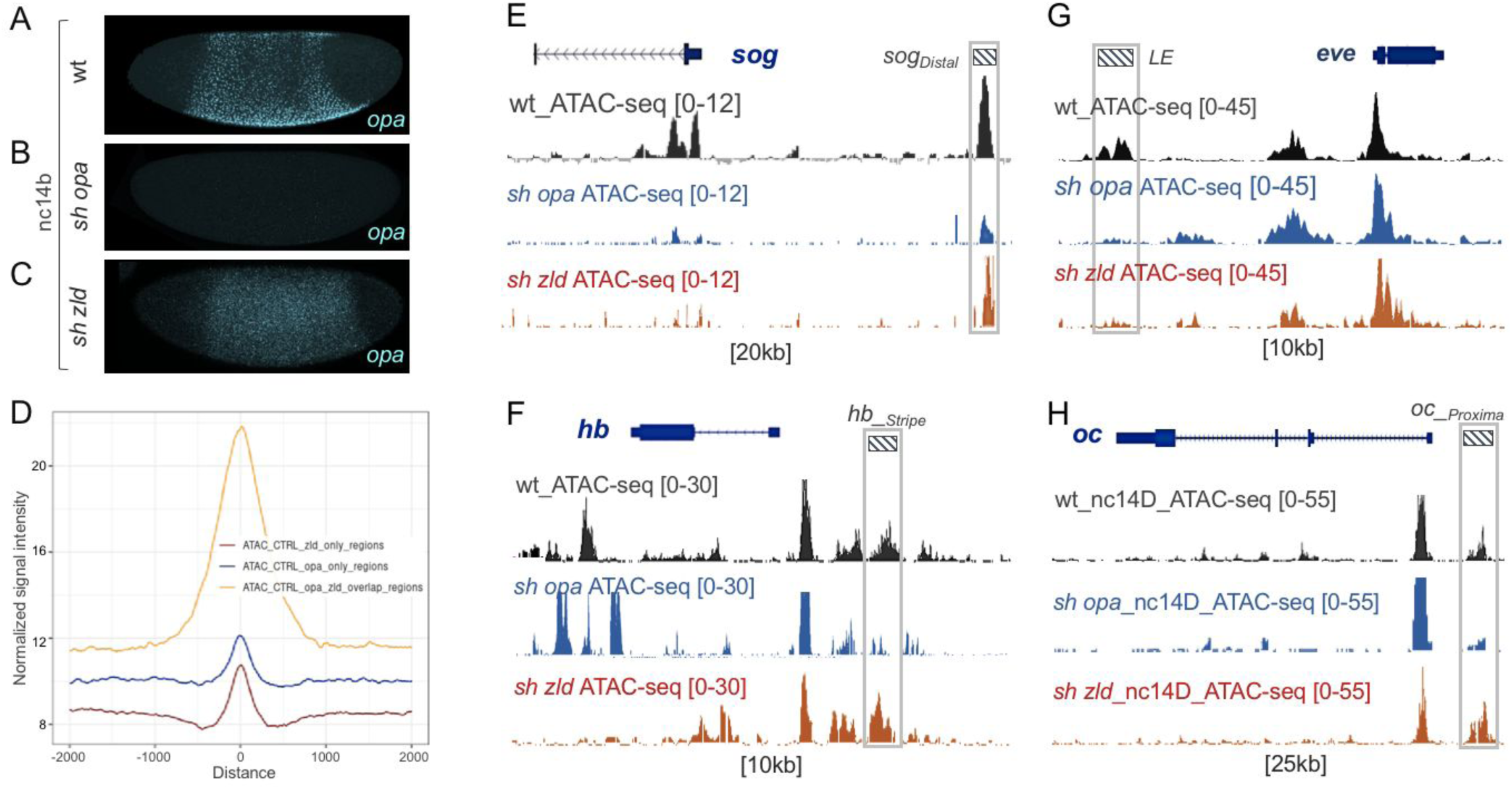
Opa is required for chromatin accessibility during MBT of the *Drosophila* embryo. (A-C) In situ hybridization using *opa* riboprobe (cyan) in either wildtype (wt; A), *opa RNAi* (*sh opa*; B), or *zld* RNAi (*sh zld*; C) embryos (n=3-5 per genotype) at stage 5. (D) Aggregation plot for wt chromatin accessibility (ATAC-Seq) signals ±2kb from the peak of Opa and Zld ChIP-seq called peaks. Signals were generated using deepTools (Ramírez et al., 2014) with RPKM method for normalization (see Methods). (E-H) UCSC dm6 genome browser tracks of representative loci showing single replicates of nc14B (E-G) and nc14D (H) ATAC-seq. Examples of late enhancer regions that significantly lose accessibility, compared to wt, either in *sh opa*-only (blue; E, F, H) or in both *sh opa* and *sh zld* (orange) (G) enhancer regions are defined by gray boxes. Plots show mean normalized read coverage of the replicates.

To provide insight into the potentially different roles of these two transcriptional factors, Opa and Zld, that support gene expression, ATAC-seq analysis was conducted on individual, single embryos either control (wt), *opa* RNAi (*sh opa*), or *zld* RNAi (*sh zld*) and the results compared. To start, we determined the relative accessibility indices for Opa-only, Zld-only, or Opa-Zld overlap regions at stage 5 (i.e. Figure 2D) using deepTools (Ramírez et al., 2014) with RPKM method for normalization (see Methods). Regions occupied by both Opa and Zld through ChIP-seq (i.e. Opa-Zld overlap) had on average 2-2.5 fold higher ATAC-seq signal (i.e. accessibility) than those that were bound by either Zld or Opa alone as detected by ChIP-seq (i.e. Zld-only or Opa-only) (Figure 4D). In summary, occupancy of both Opa and Zld is a better indicator of open chromatin regions than either factor alone. As Zld has been documented to function as a pioneer factor, that helps to make chromain accessible, these results suggest that Opa may also function in this role.

Chromatin accessibility as identified through single-embryo ATAC-seq was also examined at particular enhancers that are bound by Opa. In these Opa-bound regions, three trends were observed: (i) regions require Opa for accessibility (e.g. Figure 3E, compare with 1E; Figure 3F, compare with 3A; Figure 4 - supplemental figure 2G); (ii) regions require both Opa and Zld for accessibility (Figure 4G, compare with Figure 3B); and (iii) regions require Zld, but not Opa, for accessibility (Figure 4 - supplemental figure 1F). These results suggest that Opa can influence chromatin accessibility, but not all regions that are bound by Opa require this factor for accessibility. It is possible that at Opa-Zld overlap regions, in which both factors are bound, either Opa or Zld can suffice to support accessibility. Furthermore, in studies of Zld accessibility using a distinct method, FAIRE-seq, it was determined that, while Zld is clearly important for facilitating chromatin accessibility in the early embryo, it is not the only factor that supports this function; some chromatin regions remain accessible in *zld* mutants (Harrison et al., 2010).

### Opa-only occupied peaks require Opa to support their accessibility at mid-nc14

Co-occupancy of Opa and Zld to DNA regions correlates with increased chromatin accessibility compared to either factor alone (Figure 4D), suggesting Opa is able to influence chromatin accessibility. Furthermore, we found that in some cases Opa supports chromatin accessibility even in regions in which little or no Zld is bound (e.g. *eve_LE*; Figure 4G) and that Opa is expressed in *zld* mutants, though at reduced levels (Figure 4C). We hypothesized that Opa relates to the unidentified, Zld-independent pioneer activity.

We took a whole-genome approach towards assaying Opa’s function, and examined how chromatin accessibility is affected in Opa RNAi embryos compared to wildtype, when regions were separately analyzed based on Opa and Zld binding patterns: Opa-only, Zld-only, and Opa-Zld overlap (Figure 5A-D; also see Figure 3D). No obvious change in global accessibility was identified for the Zld-only or Opa-Zld overlap regions (Figure 5A-C); whereas, in contrast, the Opa-only regions exhibit a decreasing trend in accessibility upon Opa RNAi (*sh opa*, compare with control, wt; Figure 5D). As a control, we conducted the same type of analysis comparing wildtype to *zld* RNAi embryos, and found that all the regions were decreased upon knock-down of *zld* (*sh zld* Figure 5 - supplemental figure 1C-E). These results demonstrate that while Opa does regulate chromatin accessibility, we cannot exclude an accessory role for Zld. It is possible that these factors coordinate to regulate chromatin accessibility and/or that Zld affects Opa activity indirectly; for example, by regulating an Opa-cofactor.

**Figure 5.**
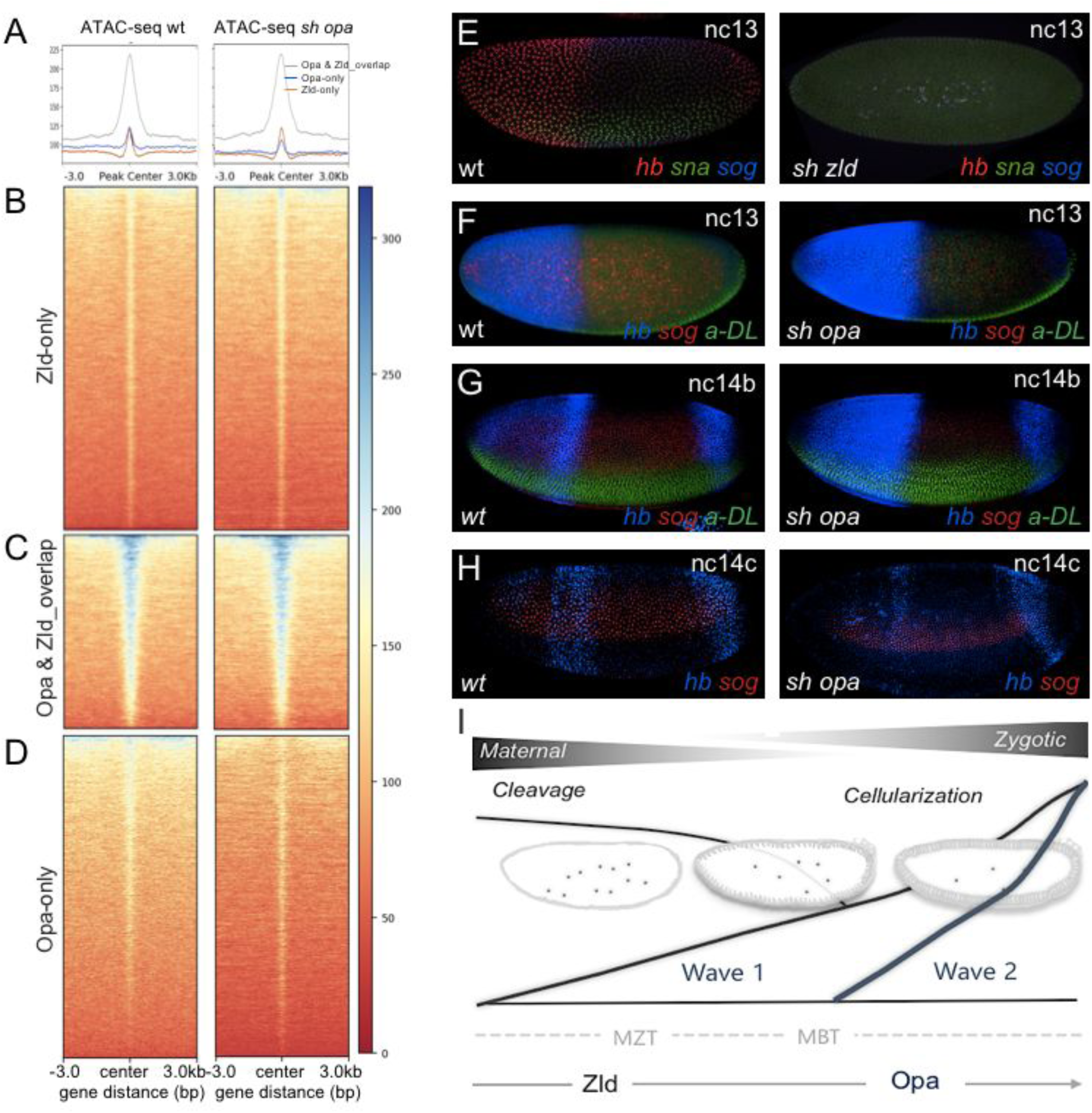
Opa is a late-acting, pioneer factor whose action follows Zelda to herald in a second wave of zygotic gene expression. **(A-D)** Aggregated signals and heatmaps of late nc14 normalized ATAC-seq signal from wt and *sh opa* mutant embryos for Zld-only (B; red trace, A), Zld &Opa overlap (C; black trace, A) and Opa-only (D; blue trace, A) ChIP-seq regions (see Figure 2D). Each row of the heatmap is a genomic region, centered to peaks of accessibility signals. The accessibility is summarized with a color code key representative of no accessibility (red) to maximum accessibility (blue). Another plot, on top of the heatmap, shows the mean signal at the genomic regions, which were centered to peaks of accessibility signals (A). **(E-G)** In situ hybridization using riboprobes to *hb* (blue) and *sog* (red), as well as anti-Dorsal staining (to highlight ventral regions) of wt and *opa RNAi* (*sh opa*) *Drosophila* embryos at indicated stages. **(I)** Schematic illustrating model supported by our results, which is that Opa, a general timing factor and likely a late-acting pioneer factor, heralds in a secondary wave of zygotic gene expression, following and coordinate with Zelda, to support the maternal-to-zygotic transition.

To determine whether these global effects on chromatin accessibility that we had observed in *opa* mutants have consequence on patterning, we examined gene expression in mutant embryos assaying for patterning phenotypes. We conducted in situ hybridizations to wildtype and *opa* RNAi embryos using riboprobes to detect endogenous transcripts for genes *sog* and *sna* expressed along the DV axis, and *hb* expressed along the AP axis. *zld* RNAi mutants were examined, in comparison; as all *sog* and *hb* genes are known to be affected in *zld* mutants, whereas *sna* is delayed in expression in *zld* mutants but recovers by nc14 (Liang et al., 2008; Nien et al., 2011). In addition, embryos were examined at two timepoints, nc13 and nc14b, corresponding to time points before and after Opa is expressed (e.g. Figure 1C). Even early, at nc13, the *zld* mutant embryos exhibit loss of expression for all three genes examined, *hb, sna*, and *sog*, supporting the view that Zld is pivotal for early expression (Figure 5E). In contrast, in *opa* mutants, at nc13, little difference was observed; *opa* is not expressed at nc13, therefore mutants would not be expected to affect patterning at this stage (Figure 5F). Later, at nc14b, the *opa* mutants do exhibit expression defects. *sog* is diminished in *opa* mutant relative to wildtype (Figure 5G); and the *hb* pattern found in *opa* mutants at nc14b is associated with an earlier pattern encompassing the full anterior cap (Koromila and Stathopoulos, 2017). It appears that the *hb_stripe* enhancer’s anterior stripe is expressed at lower levels in the *opa* mutant (Figure 5H). In contrast, the expression of *sna* in *opa* mutants appears normal (Figure 4 - supplemental figure 1D), which is the expected result as no effect on chromatin accessibility at the *sna* locus is observed (Figure 4 - supplemental figure 2A).

## DISCUSSION

In summary, our experiments indicate that Opa is a late-acting timing factor, and likely pioneer factor, which regulates gene expression across the entire embryo, along DV as well as AP axes. Previously, Opa, a non-canonical, broadly-expressed pair-rule gene, was linked only to AP axis patterning but our initial analysis of the *sog_Distal* enhancer, which is active along the DV axis, led us to investigate a more general role for this timing factor. We show that *opa* mutants also exhibit broad DV patterning changes in addition to previously identified AP patterning phenotypes. Opa likely acts directly to support this gene expression relating to patterning as Opa chromatin immunoprecipitation (ChIP) demonstrates widespread genomic binding, including at the *sog_Distal* enhancer. Additional data from single-embryo ATAC-seq provide insight into the mechanism by which Opa supports gene expression as it contributes to chromatin accessibility at a subset of regions, those occupied by Opa but not by Zld. Zld appears to predominantly support early chromatin accessibility in embryos and, subsequently, once Opa is expressed in mid-nc14 then the two factors presumably work together to support this role. However, zygotic genes (or particular enhancers) associated with mid-nc14 to gastrulation, or later, appear to be preferentially bound by Opa. Therefore, we suggest that Opa acts following the pioneer factor Zld, to also influences timing of activation at a whole-genome level through actively increasing late-acting enhancer accessibility (Figure 5G). Furthermore, Opa-mediated chromatin accessibility (Opa-only), and not just binding in general (shared by Zld), is correlated with late histone methylation. It suggests that Opa’s regulatory impact may include the control of epigenetic marks as part of its developmental program. In addition to supporting gene expression in a general manner by making chromatin accessible, Opa also presumably influences the activity of other transcription factors such as Run, co-bound to enhancers, in a mechanism that is not completely understood (Koromila and Stathopoulos, 2019; Prazak et al., 2010).

Through analysis of over-represented sites in the Opa ChIP-seq called peaks using the program HOMER, we identified a number of sites co-associated with Opa that provide additional insight into its function. For example, a motif matching the Dref consensus binding site was identified in Opa ChIP-seq peaks. Dref plays multiple roles during *Drosophila* development, as it is a multi-functional protein that has been linked to expression of signaling pathway components, being highly enriched in promoters, as well as to regulation of chromatin organization including insulator function and chromatin remodeling (rev. in Tue et al., 2017). In particular, as Dref has been linked to insulator function (Gurudatta et al., 2013) and Zld has been shown to be associated with locus-specific TAD boundary insulation (Hug et al., 2017), a future area of interest would be to examine whether temporal progression of gene regulatory networks is supported not only by the sequential action of pioneer factors that affect chromatin accessibility to affect gene activation but also affects gene expression by actively influencing chromatin conformation.

Our results also suggest that binding of Opa to DNA is enriched near promoters. Regions of accessibility are established sequentially, where enhancers are opened in advance of promoters and insulators (Blythe 2016). Opa may influence gene expression timing by affecting accessibility at promoters, possibly, in combination with Dref. Alternatively, the Trl/GAF site was associated with all classes of peaks; while present in the Opa-only and Opa-Zld overlap regions, it was enriched in the Zld-only peaks. Other studies have shown that Trl/GAF acts coordinately but separately from Zld to support chromatin accessibility in regions that do not require Zld (Moshe and Kaplan, 2017). Furthermore, Trl/GAF and Dref were shown to be associated with early versus late embryonic expression (stage 5 and later), respectively (Darbo et al., 2013; Hochheimer et al., 2002; Schulz et al., 2015). As Dref was found associated with Opa peaks but also independently linked to late embryonic expression, these results collectively support the view that Opa, like Dref, is also a late-acting factor.

More generally, the triggering of temporal waves of gene regulation in response to chromatin accessibility changes is a potentially widespread mechanism used to control developmental progression. A key future interest is to understand how the expression of genes that act as triggers, such as *opa*, are regulation. To start, the *opa* transcript is over 17 kB in length, and this relatively long transcript size may contribute to its expression in late nc14 as previously studies have suggested that transcripts of this length are difficult (or impossible) to transcribe in earlier nuclear cycles of short interphase length (i.e. nc13, etc) (Sandler et al., 2018; Shermoen and O’Farrell, 1991). *opa* expression is also regulated by the nuclear-cytoplasmic (N/C) ratio, whereas other genes like *snail* are not sensitive to N/C ratio but appear instead to be regulated by a maternal clock (Lu et al., 2009). It has been hypothesized that the initiation of genes sensitive to N/C ratio is sensitive to repression, which is diluted out with increasing N/C ratio possibly to concentrate an activator (e.g. Hamm and Harrison, 2018). We propose that genes of different timing classes may be preferentially regulated by different pioneer factors (Opa versus Zld, for example). Opa appears to be one timing factor acting at the MBT, but we propose that other factors also exist. For instance, *opa* mutants do not exhibit cellularization defects or obvious change in length of the cell cycle. It is possible that Opa regulates expression of a large number of genes at the MBT but not all genes that initiate/alter gene expression at this stage. Furthermore, *opa* expression may be regulated by maternal repression, that slowly decreases in effectiveness with dilution through increasing N/C ratio.

Opa’s expression in the embryonic trunk, but exclusion from the termini, suggest that additional late-acting factors may serve a similar role at the embryonic termini. For example, a recent study suggests that the transcription factor Orthodenticle (Otd) functions in a feed-forward relay from Bicoid to support expression of genes at the anterior of embryos (Datta et al., 2018). Furthermore, in our analysis of sites enriched in the Zld ChIP-seq dataset, we identified that the binding site for Cad transcription factor is enriched in this set. Cad is localized in a gradient that emanates from the posterior pole (Mlodzik et al., 1985). It is possible that these factors, Otd and Cad, function to support the late-activation of genes expressed at the anterior and posterior termini, possibly also functioning as pioneer factors to support action of particular late-acting enhancer in these domains where Opa is not present.

In addition, studies have shown that Zld influences morphogen gradient outputs (Foo et al., 2014), and perhaps Opa does, as well, during the secondary wave of the MZT. A late-acting morphogen gradient in the early embryo, at the time of Opa expression, is Decapentaplegic (Dpp) mediated activation of BMP signaling (Sandler and Stathopoulos, 2016; rev. in Stathopoulos and Levine, 2005b). Broadly-expressed repressors can influence temporal action of enhancers using distinct mechanisms to affect spatial gene expression patterns (e.g. Koromila and Stathopoulos, 2019), and we suggest that temporally-regulated activators such as late-acting, timing factor Opa may also support additional nuanced roles in the temporal regulation of morphogen gradient outputs. For example, our previous study showed that BMP/TGF-βtarget genes exhibit different modes of transcriptional activation, with some targets exhibiting a slower response (Sandler and Stathopoulos, 2016).

Opa is conserved as it shares homology with the Zic family of mammalian transcription factors (Aruga et al., 1996; Hursh and Stultz, 2018). Zic family members have roles in neurogenesis, myogenesis, skeletal patterning, left-right axis formation, and morphogenesis of the brain (Grinberg and Millen, 2005). In addition, Zic family members have been shown to be involved in the maintenance of pluripotency in ES cells (Lim et al., 2007). In particular, Zic3 shares significant overlap with the Oct4, Nanog, and Sox2 transcriptional networks and is important in maintaining ES cell pluripotency by preventing differentiation of cells into endodermal lineages. While we have focused on the role of Opa as an activator of gene expression, it is also possible that Opa acts to limit differentiation paths of cells.

## MATERIALS AND METHODS

### Fly stocks and husbandry

All flies were reared under standard conditions at 23°C and *yw* background was used as wild type, unless otherwise noted. *opa1/TM3,Sb* (BDGP#3212) mutant flies were rebalanced with *TM3 ftz-lacZ* marked balancer. Additionally, embryos were depleted of maternal and zygotic Opa by maternal expression of a short hairpin (*sh*) RNAi construct directed to the *opa* gene (i.e. *sh opa*). *UAS-shRNA-opa* females [TRiP.HMS01185_attP2/TM3, Sb^1^ - Bloomington *Drosophila* Stock Center (BDSC), stock #34706] were crossed to *MTD-Gal4* males (BDSC#31777). F1 *MTD-Gal4/UAS-shRNA-opa* females were crossed back to *shRNA-opa* males (#34706), and F2 embryos collected and assayed by *in situ* for *opa* mutant phenotypes. Both *sh opa* and classical *opa1* mutant embryos exhibit loss of *opa* gene expression, as detected by in situ hybridization using an *opa* riboprobe, as described (Koromila and Stathopoulos, 2017). In addition, other phenotypes of mutant embryos (i.e. *sh* hairpin and classical mutant) were compared and confirmed to be very similar to each other. Both male and female embryos were examined; sex was not determined but assumed to be equally distributed.

### Cloning

The *sog_Distal* sequences with mutated Opa or Run binding sites (i.e. *sogD_Δopa*, *sogD_Δopa4* and s*ogD_Δrun* yellow reporters) were chemically synthesized and ligated into the *eve2promoter-MS2.yellow-attB* vector using standard cloning methods as previously described (Koromila and Stathopoulos, 2019, 2017). Site-directed transgenesis was carried out using a D. melanogaster stock containing attP insertion site at position ZH-86Fb (Bloomington stock #23648). Two constructs with five and one Opa binding sites mutated (*sogD_Δopa* and *sogD_Δopa4*, respectively) were generated to check *sog_Distal’s* expression levels upon different levels of the activator Opa at nc14a and later. Mutated site sequences and their wt equivalent fragments are listed here:

- *sogD_Δrun*: tgcggtt > tAcgAtt
- *sogD_Δopa:* aaatt**cccacca** > aTGttcTcacca (1), gcgcc**cctttta** > gcgcATctttta (2), ctttt**cccacgc** > cttttcTcaTTc (3), caacg**cccgcca** > caacgcATgcca (4), gaata**cccacga** > ATGtacTcacga (5)
- *sog_DistalΔopa4*: caacg**cccgcca** > caacgcATgcca

### In situ hybridizations, immunohistochemistry, and image processing

Enzymatic in situ hybridizations were performed with antisense RNA probes labeled with digoxigenin, biotin or FITC-UTP to detect reporter or in vivo gene expression. *sna, hb, sog* (both regular and *sog* intronic), *opa* and *yellow* intronic riboprobes were used for multi-plex fluorescent in situ hybridization (FISH). For immunohistochemistry we used anti-Opa antibody (Mendoza-García et al., 2017) and anti-Zld antibody (rabbit) that is was raised for this study (1:200). Images were taken under the same settings, 26-30 Z-sections through the nuclear layer at 0.5-μm intervals, on a Zeiss LSM 880 laser-scanning microscope using a 20x air lens for fixed embryos.

### Live imaging, data acquisition and analysis

In order to monitor the various sog_Distal reporters described above in live embryos, female virgins of line yw;Nucleoporin-RFP;MCP-NoNLS-GFP were crossed with MS2 reporter-containing males. Confocal live imaging on the Zeiss LSM 880 as well as imaging optimization, segmentation, and data quantification were conducted as previously described (Koromila and Stathopoulos, 2019).

### ChIP-seq procedure

Opa-ChIP was performed as described previously (Mendoza-García et al., 2017) using chromatin prepared from 100mg of pooled collection of 3h (average of 2.5-3.5 h) and 4h (average of 3.5-4.5 h) embryos (*yw*) with anti-Opa serum. The precipitated DNA fragments were ligated with adaptors and amplified by ten cycles of PCR using NEBNext Ultra II DNAlibrary Prep Kit for Illumina (NEB) to prepare libraries for DNA sequence determination using Illumina HiSeq2500 and single-end reads of 50bp. The libraries were quantified by Qubit and Bio-Analyzer (Agilent Bioanalyzer 2100).

### ATAC-seq procedure

Embryos embryos produced from either wild-type (D. melanogaster.ZH-86Fb) or mutant *sh opa* and s*h zld* were collected on agar plates. Individual embryos were selected from plates, and nuclear morphology was observed live under a compound microscope at 20x magnification. Temperatures for sample collection were maintained at 25°C to minimize variation in cell cycle timing. Under these conditions, cell cycle times were indistinguishable. The staging of the samples started at 3-min intervals from the onset of anaphase of the previous cell cycle. Each embryo was hand-selected and hand-dechorionated for the analysis. Prepared libraries were subject to single-end sequencing, of 50bp reads, using an Illumina HiSeq2500. Fragmentation and amplification of ATAC-seq libraries were performed essentially as described previously (Blythe and Wieschaus, 2016; Buenrostro et al., 2015).

### Next generation sequencing bioinformatics

An initial analysis of the Opa-ChIP-seq data conducted by Y.I. led to the identification of the 7bp motif shown for Opa shown in Figure 1I.

The raw fastq data (50bp single-end) for Opa ChIP-seq libraries, and for wt, *sh zld* and *sh opa* ATAC-seq libraries were generated from the Illumina HiSeq2500 platform. The raw data for Zld ChIP-seq (GSM763061) and histone H3K4me1/H3K4me3 (GSE58935) were downloaded from the Gene Expression Omnibus (GEO) database. Trimmomatic-0.38 tool (Bolger et al., 2014) was used to remove Illumina adapter sequence before alignment to the *Drosophila* dm6 reference genome assembly with the Bowtie2 alignment program (Langmead and Salzberg, 2012). Alignment BAM files were subject to further sorting and duplicate removal using the Samtools package (Li et al., 2009). Reads mapped to chr2L, chr2R, chr3L, chr3R, chr4, chrX were kept and biological replicate BAM files were merged for downstream analysis. ChIP-seq and ATAC-seq signal trace files were generated using the bamCoverage function of deepTools (Ramírez et al., 2014), with RPKM r normalization and 10bp for the genomic bin size.

Both IP and input data were used for ChIP-seq peak calling. For calling of transcription factor binding sites, a workflow using bdgcmp and bdgpeakcall modules of the MACS2 peak caller (Zhang et al., 2008) was utilized. Peak calling was performed using ChIP data against input data (a proxy for genomic background). Genomic regions with q-value less than 10^-5^ were defined as ChIP-seq peak regions. Opa and Zld peak regions were combined and overlapping peaks were merged. Combined regions that overlapped both Opa and Zld peaks were defined as Opa-Zld overlap regions; regions overlapping with either Opa or Zld peaks were defined as Opa-only and Zld-only regions respectively. Further *de novo* motif analysis was performed on different ChIP-Seq regions using the HOMER program (Heinz et al., 2010) with default parameters and with options –size 200 and -mask. The most enriched *de novo* motifs identified from Opa ChIP-seq peaks and from Zld ChIP-seq peaks were queried against the Opa-Zld overlap, Opa-only and Zld-only regions for comparison and for generating aggregation plots. Average ATAC-seq signals around different ChIP-seq regions were also calculated using the annotatePeaks.pl module of HOMER, with the -size 4000 -hist 10 options used for aggregation plots. Also different ChIP-seq regions were annotated and linked to the nearest gene transcription start sites. Functional gene annotation was performed using DAVID v6.7 (https://david.ncifcrf.gov/home.jspcitation). In addition, computeMatrix and plotHeatmap modules of deepTools were used to calculate and plot normalized histone mark and ATAC-seq signal intensities around different ChIP-seq regions. DNA sequence logos were plotted using the seqLogo R package. Region overlap analysis was performed using the NeuVenn module of Neutools (doi: https://doi.org/10.1101/429639). Unless noted otherwise, R was used to generate aggregation and violin plots.

For the individual loci ATAC-seq data that are depicted in Figure 4: mapped reads were normalized similarly to a published method for better visualization (Blythe and Wieschaus, 2016): First, to define the background, 150 bp peaks were called from the original data using a very sensitive mode of HOMER (-localSize 50000-minDist 50 -size 150 -fragLength 0) to capture most of the non-background regions. These 150 bp peaks were extended from the center to form 20000 bp “signal zones”. Outside these signal zones are “background zones”. Next, to sample the background noise, 100000 150bp random regions were generated. Those 150bp random regions that completely fell into the “background zones” were regarded as “background regions”. The mean and standard deviation for the background noise were calculated from positive RPM scores of each nucleotide in these regions (ypbkg) based on log-normal distribution. Finally, RPM scores for the whole genome were centered and scaled based on the mean and standard deviation calculated, using 1 as pseudocount:

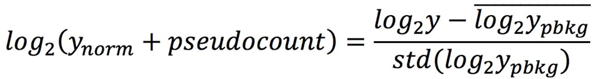

## ACKNOWLEDGEMENTS

We are grateful to Igor Antoshechkin and Henry Amrhein at the Millard and Muriel Jacobs Genetics and Genomics Laboratory at the California Institute of Technology for sequencing, the lab of Josh Dubnau for assistance with Bioanalyzer samples, David Carlson and the Institute for Advanced Computational Science at the Stony Brook University, Leslie Dunipace and Frank Macabenta for assistance with experiments and comments on the manuscript, and the entire Stathopoulos lab for helpful discussions. This study was supported by funding from NIH R35GM118146 and R03HD097535 to A.S., the Bioinformatics Resource Center at the Beckman Institute of Caltech to F.G. and L.P., and the Stony Brook University College of Arts and Sciences to J.P.G.

## AUTHOR CONTRIBUTIONS

A.S. and T.K conceived the project and planned the experimental approach. A.S. directed the project. T.K performed wet experiments except ChIP-seq, which was performed by Y.I. with support of the Caltech genomics core. The computational approach was overseen by F.G., P.H., and T.K.; F.G. performed all computational analysis except normalization of ATAC-seq data for visualization of individual loci, which was performed by P.H. An initial, independent analysis of the Opa-ChIP-seq data was conducted by Y.I. Quantitative analysis of imaging data was performed by T.K. Data were analyzed by A.S., T.K., and F.G. The manuscript was written by A.S. and T.K. with input and editing help from F.G., Y.I, P.H., L.P. and J.P.G.

## COMPETING INTERESTS

No competing interests declared.

## DATA AND CODE AVAILABILITY

GEO accession numbers: ChIP-seq and ATAC-seq (GSE140722).

The codes for Opa ChIP-seq and ATAC-seq processing (alignment and peak calling) were uploaded to github: https://github.com/gaofan83/Stathopoulos_Lab

## REFERENCES

Abdusselamoglu MD, Eroglu E, Burkard TR, Knoblich JA. 2019. The transcription factor odd-paired regulates temporal identity in transit-amplifying neural progenitors via an incoherent feed-forward loop. Elife 8. doi:10.7554/eLife.46566

Ali-Murthy Z, Kornberg TB. 2016. Bicoid gradient formation and function in the Drosophila pre-syncytial blastoderm. Elife 5. doi:10.7554/eLife.13222

Aruga J, Millen KJ. 2018. ZIC1 Function in Normal Cerebellar Development and Human Developmental Pathology. Adv Exp Med Biol 1046:249–268.

Aruga J, Nagai T, Tokuyama T, Hayashizaki Y, Okazaki Y, Chapman VM, Mikoshiba K. 1996. The mouse zic gene family. Homologues of the Drosophila pair-rule gene odd-paired. J Biol Chem 271:1043–1047.

Benedyk MJ, Mullen JR, DiNardo S. 1994. odd-paired: a zinc finger pair-rule protein required for the timely activation of engrailed and wingless in Drosophila embryos. Genes Dev 8:105–117.

Blythe SA, Wieschaus EF. 2016. Establishment and maintenance of heritable chromatin structure during early embryogenesis. Elife 5. doi:10.7554/eLife.20148

Bolger AM, Lohse M, Usadel B. 2014. Trimmomatic: a flexible trimmer for Illumina sequence data. Bioinformatics 30:2114–2120.

Bonn S, Zinzen RP, Girardot C, Gustafson EH, Perez-Gonzalez A, Delhomme N, Ghavi-Helm Y, Wilczyński B, Riddell A, Furlong EEM. 2012. Tissue-specific analysis of chromatin state identifies temporal signatures of enhancer activity during embryonic development. Nat Genet 44:148–156.

Bothma JP, Garcia HG, Esposito E, Schlissel G, Gregor T, Levine M. 2014. Dynamic regulation of eve stripe 2 expression reveals transcriptional bursts in living Drosophila embryos. Proc Natl Acad Sci U S A 111:10598–10603.

Buenrostro JD, Wu B, Litzenburger UM, Ruff D, Gonzales ML, Snyder MP, Chang HY, Greenleaf WJ. 2015. Single-cell chromatin accessibility reveals principles of regulatory variation. Nature 523:486–490.

Cadigan KM, Grossniklaus U, Gehring WJ. 1994. Localized expression of sloppy paired protein maintains the polarity of Drosophila parasegments. Genes Dev 8:899–913.

Chen H, Xu Z, Mei C, Yu D, Small S. 2012. A System of Repressor Gradients Spatially Organizes the Boundaries of Bicoid-Dependent Target Genes. Cell. doi:10.1016/j.cell.2012.03.018

Chen K, Johnston J, Shao W, Meier S, Staber C, Zeitlinger J. 2013. A global change in RNA polymerase II pausing during the Drosophila midblastula transition. Elife 2:e00861.

Cimbora DM, Sakonju S. 1995. Drosophila midgut morphogenesis requires the function of the segmentation gene odd-paired. Dev Biol 169:580–595.

Clark E, Akam M. 2016. Odd-paired controls frequency doubling in segmentation by altering the pair-rule gene regulatory network. Elife 5. doi:10.7554/eLife.18215

Clark E, Peel AD. 2018. Evidence for the temporal regulation of insect segmentation by a conserved sequence of transcription factors. Development. doi:10.1242/dev.155580

Darbo E, Herrmann C, Lecuit T, Thieffry D, van Helden J. 2013. Transcriptional and epigenetic signatures of zygotic genome activation during early Drosophila embryogenesis. BMC Genomics 14:226.

Datta RR, Ling J, Kurland J, Ren X, Xu Z, Yucel G, Moore J, Shokri L, Baker I, Bishop T, Struffi P, Levina R, Bulyk ML, Johnston RJ Jr, Small S. 2018. A feed-forward relay integrates the regulatory activities of Bicoid and Orthodenticle via sequential binding to suboptimal sites. Genes Dev 32:723–736.

Foe VE, Alberts BM. 1983. Studies of nuclear and cytoplasmic behaviour during the five mitotic cycles that precede gastrulation in Drosophila embryogenesis. J Cell Sci 61:31–70.

Foo SM, Sun Y, Lim B, Ziukaite R, O’Brien K, Nien C-Y, Kirov N, Shvartsman SY, Rushlow CA. 2014. Zelda potentiates morphogen activity by increasing chromatin accessibility. Curr Biol 24:1341–1346.

Fujioka M, Miskiewicz P, Raj L, Gulledge AA, Weir M, Goto T. 1996. Drosophila Paired regulates late even-skipped expression through a composite binding site for the paired domain and the homeodomain. Development 122:2697–2707.

Grinberg I, Millen KJ. 2005. The ZIC gene family in development and disease. Clin Genet 67:290–296.

Gurudatta BV, Yang J, Van Bortle K, Donlin-Asp PG, Corces VG. 2013. Dynamic changes in the genomic localization of DNA replication-related element binding factor during the cell cycle. Cell Cycle 12:1605–1615.

Hamm DC, Harrison MM. 2018. Regulatory principles governing the maternal-to-zygotic transition: insights from Drosophila melanogaster. Open Biol 8:180183.

Hang S, Gergen JP. 2017. Different modes of enhancer-specific regulation by Runt and Even-skipped during segmentation. Mol Biol Cell 28:681–691.

Harrison MM, Botchan MR, Cline TW. 2010. Grainyhead and Zelda compete for binding to the promoters of the earliest-expressed Drosophila genes. Dev Biol 345:248–255.

Harrison MM, Li X-Y, Kaplan T, Botchan MR, Eisen MB. 2011. Zelda binding in the early Drosophila melanogaster embryo marks regions subsequently activated at the maternal-to-zygotic transition. PLoS Genet 7:e1002266.

Heinz S, Benner C, Spann N, Bertolino E, Lin YC, Laslo P, Cheng JX, Murre C, Singh H, Glass CK. 2010. Simple combinations of lineage-determining transcription factors prime cis-regulatory elements required for macrophage and B cell identities. Mol Cell 38:576–589.

Hirono K, Margolis JS, Posakony JW, Doe CQ. 2012. Identification of hunchback cis-regulatory DNA conferring temporal expression in neuroblasts and neurons. Gene Expr Patterns 12:11–17.

Hirose F, Yamaguchi M, Handa H, Inomata Y, Matsukage A. 1993. Novel 8-base pair sequence (Drosophila DNA replication-related element) and specific binding factor involved in the expression of Drosophila genes for DNA polymerase alpha and proliferating cell nuclear antigen. J Biol Chem 268:2092–2099.

Hochheimer A, Zhou S, Zheng S, Holmes MC, Tjian R. 2002. TRF2 associates with DREF and directs promoter-selective gene expression in Drosophila. Nature 420:439–445.

Hong J-W, Hendrix DA, Levine MS. 2008. Shadow enhancers as a source of evolutionary novelty. Science 321:1314.

Houtmeyers R, Souopgui J, Tejpar S, Arkell R. 2013. The ZIC gene family encodes multi-functional proteins essential for patterning and morphogenesis. Cell Mol Life Sci 70:3791–3811.

Hug CB, Grimaldi AG, Kruse K, Vaquerizas JM. 2017. Chromatin Architecture Emerges during Zygotic Genome Activation Independent of Transcription. Cell 169:216–228.e19.

Hursh DA, Stultz BG. 2018. Odd-Paired: The Drosophila Zic Gene. Adv Exp Med Biol 1046:41–58.

Ip YT, Park RE, Kosman D, Bier E, Levine M. 1992. The dorsal gradient morphogen regulates stripes of rhomboid expression in the presumptive neuroectoderm of the Drosophila embryo. Genes Dev 6:1728–1739.

Iwafuchi-Doi M, Zaret KS. 2014. Pioneer transcription factors in cell reprogramming. Genes Dev 28:2679–2692.

Khan A, Fornes O, Stigliani A, Gheorghe M, Castro-Mondragon JA, van der Lee R, Bessy A, Chèneby J, Kulkarni SR, Tan G, Baranasic D, Arenillas DJ, Sandelin A, Vandepoele K, Lenhard B, Ballester B, Wasserman WW, Parcy F, Mathelier A. 2018. JASPAR 2018: update of the open-access database of transcription factor binding profiles and its web framework. Nucleic Acids Res 46:D1284.

Koenecke N, Johnston J, He Q, Meier S, Zeitlinger J. 2017. Drosophila poised enhancers are generated during tissue patterning with the help of repression. Genome Res 27:64–74.

Koromila T, Stathopoulos A. 2019. Distinct Roles of Broadly Expressed Repressors Support Dynamic Enhancer Action and Change in Time. Cell Rep 28:855–863.e5.

Koromila T, Stathopoulos A. 2017. Broadly expressed repressors integrate patterning across orthogonal axes in embryos. Proc Natl Acad Sci U S A. doi:10.1073/pnas.1703001114

Kvon EZ, Kazmar T, Stampfel G, Omar Yáñez-Cuna J, Pagani M, Schernhuber K, Dickson BJ, Stark A. 2014. Genome-scale functional characterization of Drosophila developmental enhancers in vivo. Nature. doi:10.1038/nature13395

Langmead B, Salzberg SL. 2012. Fast gapped-read alignment with Bowtie 2. Nat Methods 9:357–359.

Leichsenring M, Maes J, Mössner R, Driever W, Onichtchouk D. 2013. Pou5f1 transcription factor controls zygotic gene activation in vertebrates. Science 341:1005–1009.

Liang H-L, Nien C-Y, Liu H-Y, Metzstein MM, Kirov N, Rushlow C. 2008. The zinc-finger protein Zelda is a key activator of the early zygotic genome in Drosophila. Nature 456:400–403.

Li H, Handsaker B, Wysoker A, Fennell T, Ruan J, Homer N, Marth G, Abecasis G, Durbin R, 1000 Genome Project Data Processing Subgroup. 2009. The Sequence Alignment/Map format and SAMtools. Bioinformatics 25:2078–2079.

Lim LS, Hong FH, Kunarso G, Stanton LW. 2010. The pluripotency regulator Zic3 is a direct activator of the Nanog promoter in ESCs. Stem Cells 28:1961–1969.

Lim LS, Loh Y-H, Zhang W, Li Y, Chen X, Wang Y, Bakre M, Ng H-H, Stanton LW. 2007. Zic3 Is Required for Maintenance of Pluripotency in Embryonic Stem Cells. Molecular Biology of the Cell. doi:10.1091/mbc.e06-07-0624

Li X-Y, Harrison MM, Villalta JE, Kaplan T, Eisen MB. 2014. Establishment of regions of genomic activity during the Drosophila maternal to zygotic transition. eLife. doi:10.7554/elife.03737

Lott SE, Villalta JE, Schroth GP, Luo S, Tonkin LA, Eisen MB. 2011. Noncanonical compensation of zygotic X transcription in early Drosophila melanogaster development revealed through single-embryo RNA-seq. PLoS Biol 9:e1000590.

Lucas T, Tran H, Perez Romero CA, Guillou A, Fradin C, Coppey M, Walczak AM, Dostatni N. 2018. 3 minutes to precisely measure morphogen concentration. PLoS Genet 14:e1007676.

Lu X, Li JM, Elemento O, Tavazoie S, Wieschaus EF. 2009. Coupling of zygotic transcription to mitotic control at the Drosophila mid-blastula transition. Development 136:2101–2110.

Markstein M, Markstein P, Markstein V, Levine MS. 2002. Genome-wide analysis of clustered Dorsal binding sites identifies putative target genes in the Drosophila embryo. Proc Natl Acad Sci U S A 99:763–768.

McDaniel SL, Gibson TJ, Schulz KN, Fernandez Garcia M, Nevil M, Jain SU, Lewis PW, Zaret KS, Harrison MM. 2019. Continued Activity of the Pioneer Factor Zelda Is Required to Drive Zygotic Genome Activation. Mol Cell 74:185–195.e4.

Mendoza-García P, Hugosson F, Fallah M, Higgins ML, Iwasaki Y, Pfeifer K, Wolfstetter G, Varshney G, Popichenko D, Gergen JP, Hens K, Deplancke B, Palmer RH. 2017. The Zic family homologue Odd-paired regulates Alk expression in Drosophila. PLoS Genet 13:e1006617.

Mlodzik M, Fjose A, Gehring WJ. 1985. Isolation of caudal, a Drosophila homeo box-containing gene with maternal expression, whose transcripts form a concentration gradient at the pre-blastoderm stage. The EMBO Journal. doi:10.1002/j.1460-2075.1985.tb04030.x

Moshe A, Kaplan T. 2017. Genome-wide search for Zelda-like chromatin signatures identifies GAF as a pioneer factor in early fly development. Epigenetics Chromatin 10:33.

Nien C-Y, Liang H-L, Butcher S, Sun Y, Fu S, Gocha T, Kirov N, Manak JR, Rushlow C. 2011. Temporal coordination of gene networks by Zelda in the early Drosophila embryo. PLoS Genet 7:e1002339.

Ohler U, Liao G-C, Niemann H, Rubin GM. 2002. Computational analysis of core promoters in the Drosophila genome. Genome Biol 3:RESEARCH0087.

Ozdemir A, Fisher-Aylor KI, Pepke S, Samanta M, Dunipace L, McCue K, Zeng L, Ogawa N, Wold BJ, Stathopoulos A. 2011. High resolution mapping of Twist to DNA in Drosophila embryos: Efficient functional analysis and evolutionary conservation. Genome Res 21:566–577.

Pearson JC, Crews ST. 2014. Enhancer diversity and the control of a simple pattern of Drosophila CNS midline cell expression. Dev Biol 392:466–482.

Perry MW, Boettiger AN, Levine M. 2011. Multiple enhancers ensure precision of gap gene-expression patterns in the Drosophila embryo. Proc Natl Acad Sci U S A 108:13570–13575.

Perry MW, Bothma JP, Luu RD, Levine M. 2012. Precision of hunchback expression in the Drosophila embryo. Curr Biol 22:2247–2252.

Prazak L, Fujioka M, Gergen JP. 2010. Non-additive interactions involving two distinct elements mediate sloppy-paired regulation by pair-rule transcription factors. Dev Biol 344:1048–1059.

Rada-Iglesias A, Bajpai R, Swigut T, Brugmann SA, Flynn RA, Wysocka J. 2011. A unique chromatin signature uncovers early developmental enhancers in humans. Nature 470:279–283.

Ramírez F, Dündar F, Diehl S, Grüning BA, Manke T. 2014. deepTools: a flexible platform for exploring deep-sequencing data. Nucleic Acids Res 42:W187–91.

Rogers WA, Goyal Y, Yamaya K, Shvartsman SY, Levine MS. 2017. Uncoupling neurogenic gene networks in the embryo. Genes Dev 31:634–638.

Sandler JE, Irizarry J, Stepanik V, Dunipace L, Amrhein H, Stathopoulos A. 2018. A Developmental Program Truncates Long Transcripts to Temporally Regulate Cell Signaling. Dev Cell 47:773–784.e6.

Sandler JE, Stathopoulos A. 2016. Quantitative Single-Embryo Profile of Drosophila Genome Activation and the Dorsal-Ventral Patterning Network. Genetics 202:1575–1584.

Schulz KN, Bondra ER, Moshe A, Villalta JE, Lieb JD, Kaplan T, McKay DJ, Harrison MM. 2015. Zelda is differentially required for chromatin accessibility, transcription factor binding, and gene expression in the early Drosophila embryo. Genome Res 25:1715–1726.

Schulz KN, Harrison MM. 2019. Mechanisms regulating zygotic genome activation. Nat Rev Genet 20:221–234.

Sen A, Stultz BG, Lee H, Hursh DA. 2010. Odd paired transcriptional activation of decapentaplegic in the Drosophila eye/antennal disc is cell autonomous but indirect. Dev Biol 343:167–177.

Shermoen AW, McCleland ML, O’Farrell PH. 2010. Developmental Control of Late Replication and S Phase Length. Current Biology. doi:10.1016/j.cub.2010.10.021

Shermoen AW, O’Farrell PH. 1991. Progression of the cell cycle through mitosis leads to abortion of nascent transcripts. Cell. doi:10.1016/0092-8674(91)90182-x

Shindo Y, Amodeo AA. 2019. Dynamics of Free and Chromatin-Bound Histone H3 during Early Embryogenesis. Curr Biol 29:359–366.e4.

Shir-Shapira H, Sharabany J, Filderman M, Ideses D, Ovadia-Shochat A, Mannervik M, Juven-Gershon T. 2015. Structure-function analysis of the Drosophila melanogaster Caudal transcription factor provides insights into core promoter-preferential activation. J Biol Chem 290:20747.

Staller MV, Yan D, Randklev S, Bragdon MD, Wunderlich ZB, Tao R, Perkins LA, Depace AH, Perrimon N. 2013. Depleting gene activities in early Drosophila embryos with the “maternal-Gal4-shRNA” system. Genetics 193:51–61.

Stathopoulos A, Levine M. 2005a. Localized repressors delineate the neurogenic ectoderm in the early Drosophila embryo. Dev Biol 280:482–493.

Stathopoulos A, Levine M. 2005b. Genomic regulatory networks and animal development. Dev Cell 9:449–462.

Swantek D, Gergen JP. 2004. Ftz modulates Runt-dependent activation and repression of segment-polarity gene transcription. Development 131:2281–2290.

Tadros W, Lipshitz HD. 2009. The maternal-to-zygotic transition: a play in two acts. Development 136:3033–3042.

Tue NT, Yoshioka Y, Mizoguchi M, Yoshida H, Zurita M, Yamaguchi M. 2017. DREF plays multiple roles during Drosophila development. Biochim Biophys Acta Gene Regul Mech 1860:705–712.

Vastenhouw NL, Cao WX, Lipshitz HD. 2019. The maternal-to-zygotic transition revisited. Development 146. doi:10.1242/dev.161471

Xu Z, Chen H, Ling J, Yu D, Struffi P, Small S. 2014. Impacts of the ubiquitous factor Zelda on Bicoid-dependent DNA binding and transcription in Drosophila. Genes Dev 28:608–621.

Yamada S, Whitney PH, Huang S-K, Eck EC, Garcia HG, Rushlow CA. 2019. The Drosophila Pioneer Factor Zelda Modulates the Nuclear Microenvironment of a Dorsal Target Enhancer to Potentiate Transcriptional Output. Curr Biol 29:1387–1393.e5.

Yuan K, Seller CA, Shermoen AW, O’Farrell PH. 2016. Timing the Drosophila Mid-Blastula Transition: A Cell Cycle-Centered View. Trends Genet 32:496–507.

Zhang Y, Liu T, Meyer CA, Eeckhoute J, Johnson DS, Bernstein BE, Nusbaum C, Myers RM, Brown M, Li W, Liu XS. 2008. Model-based analysis of ChIP-Seq (MACS). Genome Biol 9:R137.

