## Supplemental Material for "Odd-paired is a late-acting pioneer factor coordinating with Zelda to broadly regulate gene expression in early embryos"

**Figure 1 - figure supplement 1**

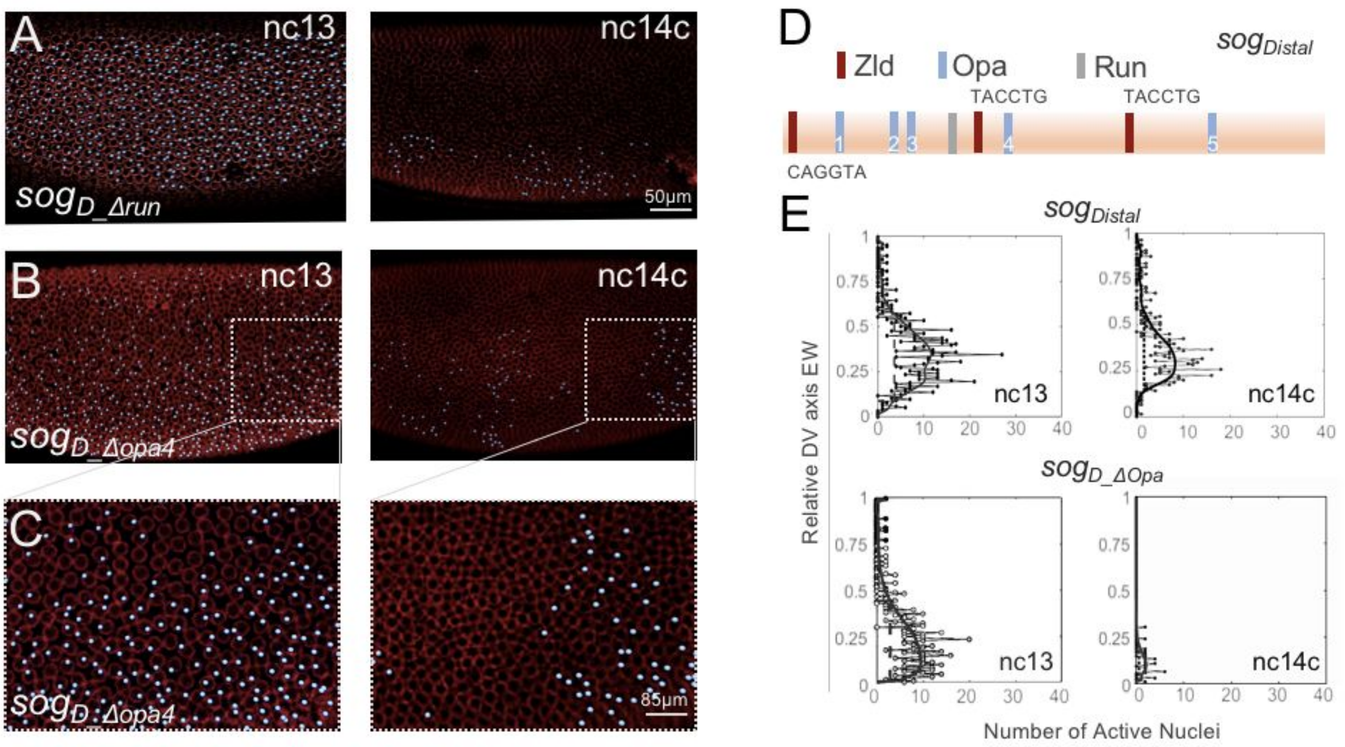


**Figure 1 - figure supplement 1. Assay of *sog_Distal* expression outputs through live in vivo imaging following mutagenesis of Opa or Run predicted binding sites.**

(A-C) MS2-MCP imaging-based computationally defined dots of nascent transcripts (blue) and nuclear membranes marked by Nup-RFP (ed) associated with the $sog_D$*_Δrun* reporter in which the single Run site (see D) is mutated (Koromila and Stathopoulos, 2019) or the $sog_D$*_Δopa4* reporter, in which only one Opa binding site, site 4 (D), is mutated. (C) shows magnified views of B; expression is sometimes retained at the posterior of embryos.

(D) 650 bp *sog_Distal* enhancer sequence showing relative organization of binding sites for transcription factors based on matches to consensus sequences: 3 sites for Zld, 5 sites for Opa, and 1 site for Run. Base-pair sequences for three Zld sites shown, which are direct matches to the consensus. Zld and Run sites were previously mutated and characterized (Foo et al., 2014; Koromila and Stathopoulos, 2019).

(E) Plots of number of active nuclei, defined by counting dots (x-axis) versus relative DV axis embryo-width (EW) position (y-axis), as analyzed for representative stills from nc13 and nc14c. Black dotted traces represent raw counts of active nuclei-dots of MS2-MCP signal (bins represent minimum of four dots) detected throughout embryos containing indicated constructs after projection of scans of particular time points were collapsed along the anterior-posterior (AP) axis and dots counted and binned across the DV axis (EW) (for details see: Koromila and Stathopoulos, 2019). The black line for either *sogDistal* wild-type or mutant construct represents normalization after applying a smoothing curve.

**Figure 1 - supplemental figure 2**

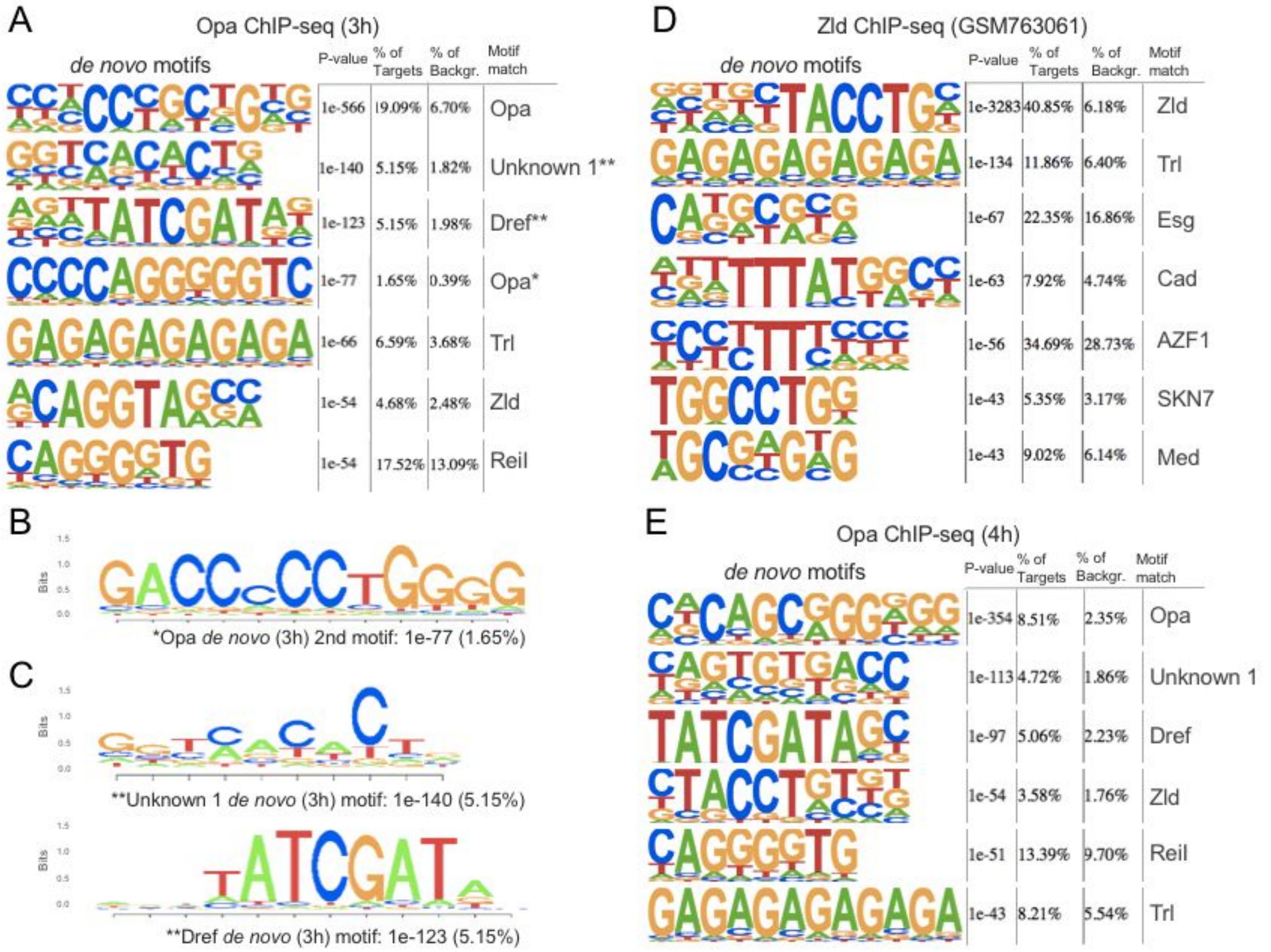


**Figure 1 - supplemental figure 2. Most abundant motifs identified using HOMER *de novo* motif analysis within the two Opa ChIP-seq and Zld ChIP-seq datasets.**

**(A)** Seven most enriched motifs for Opa (3h) ChIP-seq and Zld (stage 5) ChIP-seq analysis as detected by HOMER. The top identified motifs also shown in Figure 1. A second motif exhibiting extended homology with the JASPAR Opa consensus was identified at lower frequency (see panel B).

**(B)** Reverse complement of fourth most abundant motifs identified using HOMER within the 3h Opa ChIP-seq dataset, showing extended homology to JASPAR Opa site.

**(C)** Opa ChiP-seq six most enriched motifs for 4h embryos. The top identified motif shown also in Figure 1J.

**Figure 2 - supplemental table 1**

| | I. Opa-only | II. Zld-only | III. Opa-Zld overlap |
|---|---|---|---|
| **Opa-like** | **1e-522 (15.97%)** | n.f. | **1e-43 (4.54%)** |
| **Zelda-like** | n.f. | **1e-2833 (55.03%)** | **1e-240 (15.05%)** |
| **Trl-like** | 1e-12 (6.78%) | 1e-23 (4.45%) | **1e-76 (10.56%)** |
| **Dref-like** | **1e-119 (6.08%)** | n.f | 1e-19 (4.13%) |
| **Cad-like** | n.f. | **1e-63 (14.18%)** | n.f. |
| **Unknown 1** | **1e-117 (5.93%)** | n.f. | **1e-33 (2.99%)** |
| **Run-like** | 1e-22 (2.51%) | n.f. | n.f. |

**Table 1. Abundance of motifs for known transcription factors found by HOMER within the three corresponding sets of peaks.** Each set of peaks (i.e. Opa-only, Zld-only, and Opa-Zld overlap) was independently analyzed by HOMER, and binding site consensus sequences defined independently for the listed transcription factors. If no motif for a certain transcription was designated significant by HOMER within a particular set of peaks, designed not found (n.f.). P-values are shown for enrichment of consensus binding sites within each class; abundance shown in parentheses. Data for a subset of sites are shown in Figure 2.

**Figure 2 -supplemental figure 1**

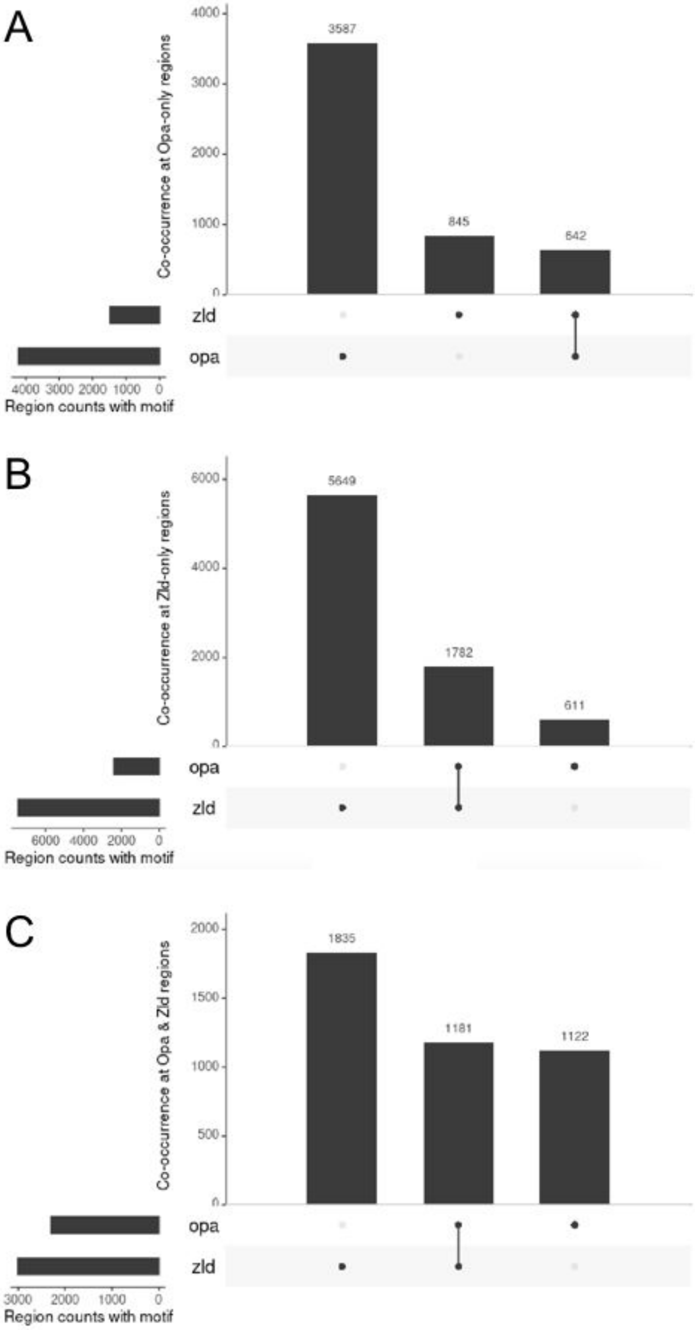


**Figure 2 - supplemental figure 1. Overlap statistics of regions containing Opa and Zld motifs: a novel visualization technique for the quantitative analysis of sets, their intersections, and aggregates of intersections.** (A-C) The UpSet plots showing enrichment of top *de novo* motifs identified from Opa (3h) and Zld (GSM763061) peaks. Motif scanning was performed within ±500bp of the region center.

Each column corresponds to an Opa/Zld set as indicated in Figure 2A-C, and each row corresponds to one segment in the Venn diagram (Figure 2D). Opa-only peaks (A); Zld-only peaks (B); Opa-Zld overlap peaks (C).

**Figure 3 - supplemental figure 1**

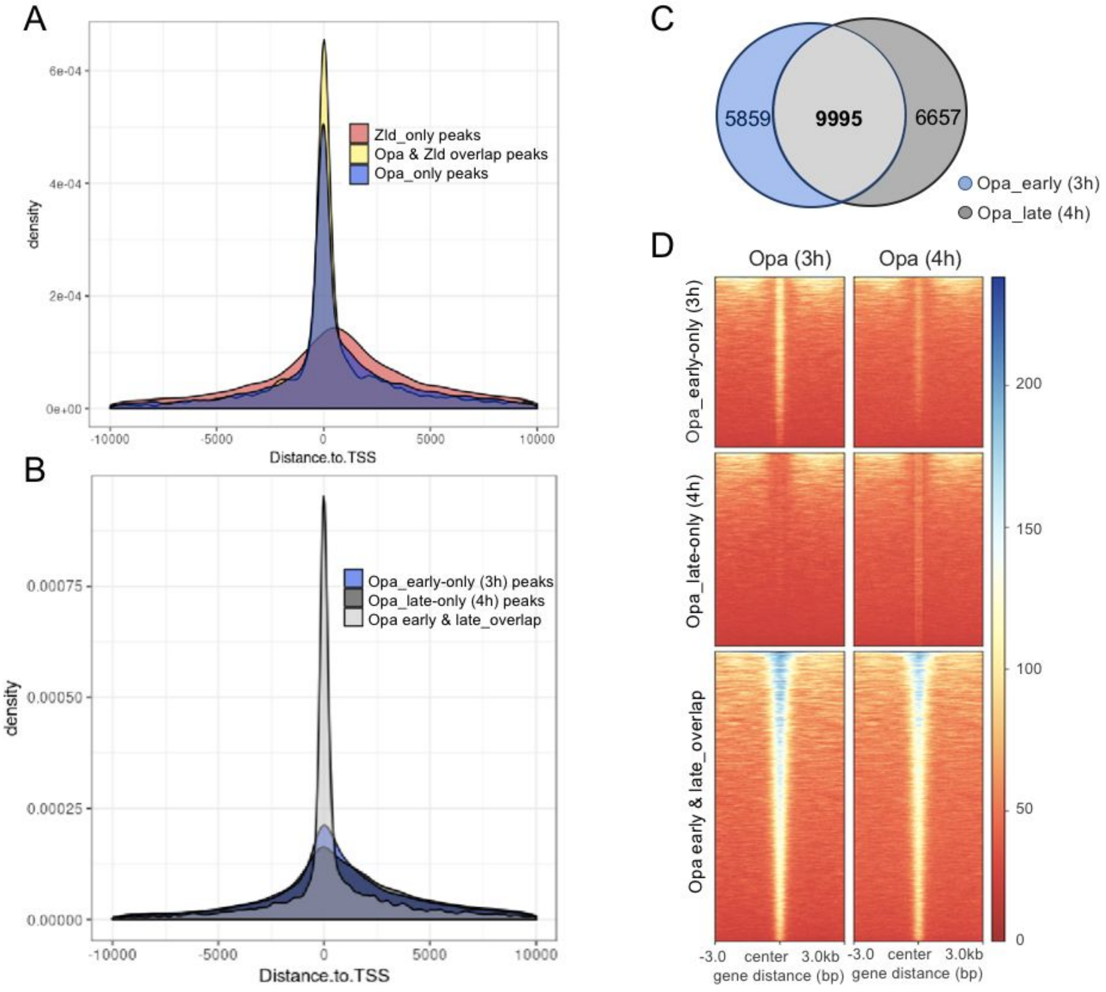


**Figure 3 - supplemental figure 1. Position of peaks relative to transcription start site as well as a comparison of Opa_early (3h) versus Opa_late (4h) peak size and overlap.**

(A,B) Histograms showing the distribution of the position of Opa-only, Opa_Zld overlap and Zld only peaks (A) or Opa_early only, Opa_late only, and Opa early & late overlap peaks (B) relative to the transcription start site (TSS).

(C) Venn diagram comparing number of Opa_early (3h) versus Opa_late (4h) ChIP-seq called peaks.

(D) Heatmaps of normalized ChIP-seq data of Opa-early (3h; left) or Opa-late (4h; right) centered on genomic sequences representing called peaks of one of three classes: Opa_early only (3h), Opa_late only

(4h), or Opa early & late overlap. Center of ChIP-seq called peaks are positioned at 0, and extended to 3 kb shown on either side. The size of the peaks is displayed as a color code from blue (largest) to red (smallest). Opa_early only (3h; top), Opa_late only (4h; right) ChIP-seq regions split up to illustrate relative size of peaks and enrichment of these DNA regions in the two ChIP-seq samples. Key indicates normalized signal intensities (see Methods) around different ChIP-seq regions.

**Figure 4 - supplemental figure 1**

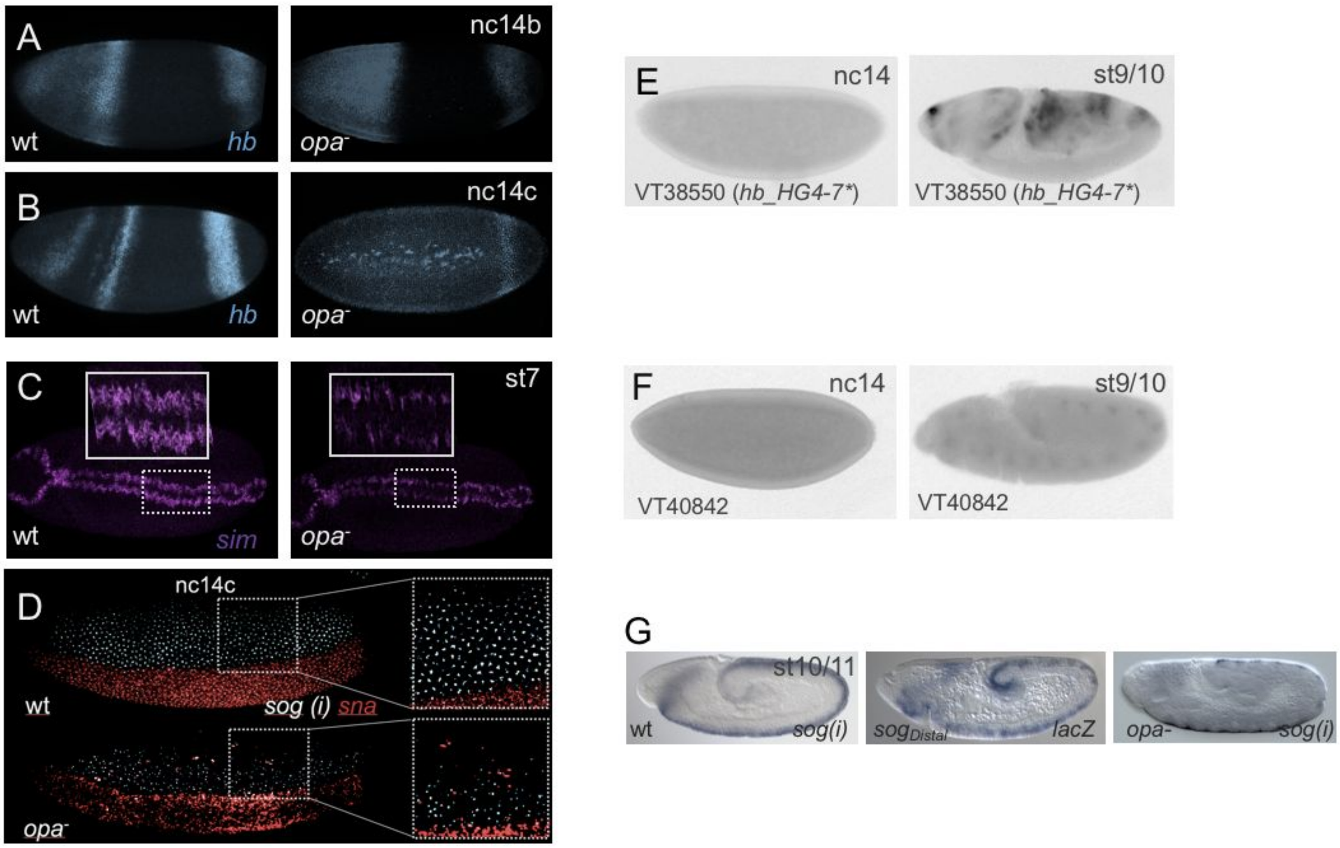


**Figure 4 - supplemental figure 1. *opa1* mutants exhibit broad patterning defects affecting DV as well as AP patterning.** In situ hybridization using riboprobes to detect transcripts for *hb* (blue; A,B), *sim* (purple; C), *sna* (red; D), intronic *sog* [*sog(i)*] (blue; D), and *lacZ* (E,F,G) or *sog(i)* (G) in *Drosophila* embryos at the indicated stages.

(A,B) *hb* expression in wt embryos compared to *opa*$^1$ mutant embryos at two stages: nc14b and nc14c.

(C) *sim* expression in wt embryos compared to *opa*$^1$ mutant embryos at stage 7.

(D) *sog(i)* and *sna* expression in wt embryos compared to *opa*$^1$ mutant embryos at nc14c.

(E,F) Data from publicly available genome-scale enhancer characterization (Kvon et al., 2014) demonstrating late expression (stage 9/10) for *hb_GH4-7* enhancer associated with *hb* locus and VT40842 enhancer associated with *sim* locus.

(G) *sog(i)* expression in wt embryo ast stage 10/11 (left) is comparable to expression supported by the *sog_Distal* enhancer in embryos (middle) at this same stage; whereas *sog(i)* expression is diminished in *opa*$^{1}$ mutant embryo (right).

**Figure 4 - supplemental figure 2**

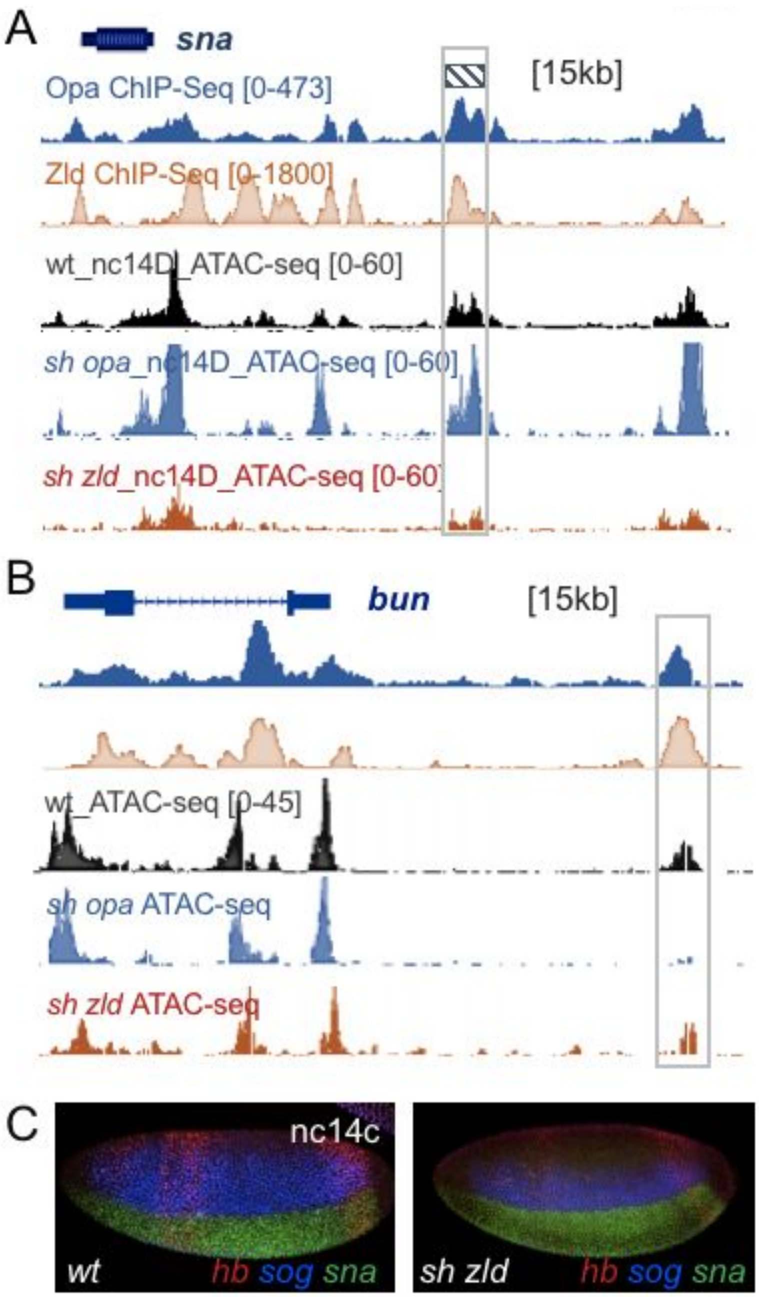


**Figure 4 - supplemental figure 2. Additional examples of ATAC-seq data showing differences in chromatin accessibility in response to Opa or Zld RNAi.**

(A,B) UCSC dm6 genome browser tracks of representative loci showing single replicates of stage 5 single embryo normalized ATAC-seq data in comparison to Opa and Zld ChIP-seq (stage 5). (A) Examples of an enhancer region, associated with gene *snail* (*sna*) that exhibits decreased accessibility in *sh zld*, compared to wt, but exhibit no decrease in *sh opa* at nc14D. (B) Another example of an enhancer region associated with the gene *bunched* (*bun*) that exhibits decreased accessibility in *sh opa* but little change in *sh zld* at nc14B. Enhancer regions of focus are defined by gray boxes.

(C) In situ hybridization using riboprobes to detect transcripts for *hb* (red), *sog* (blue), *sna* (red; D), in *Drosophila* embryos, wt or *sh zld*, at nc14c.

**Figure 5 - supplemental figure 1**

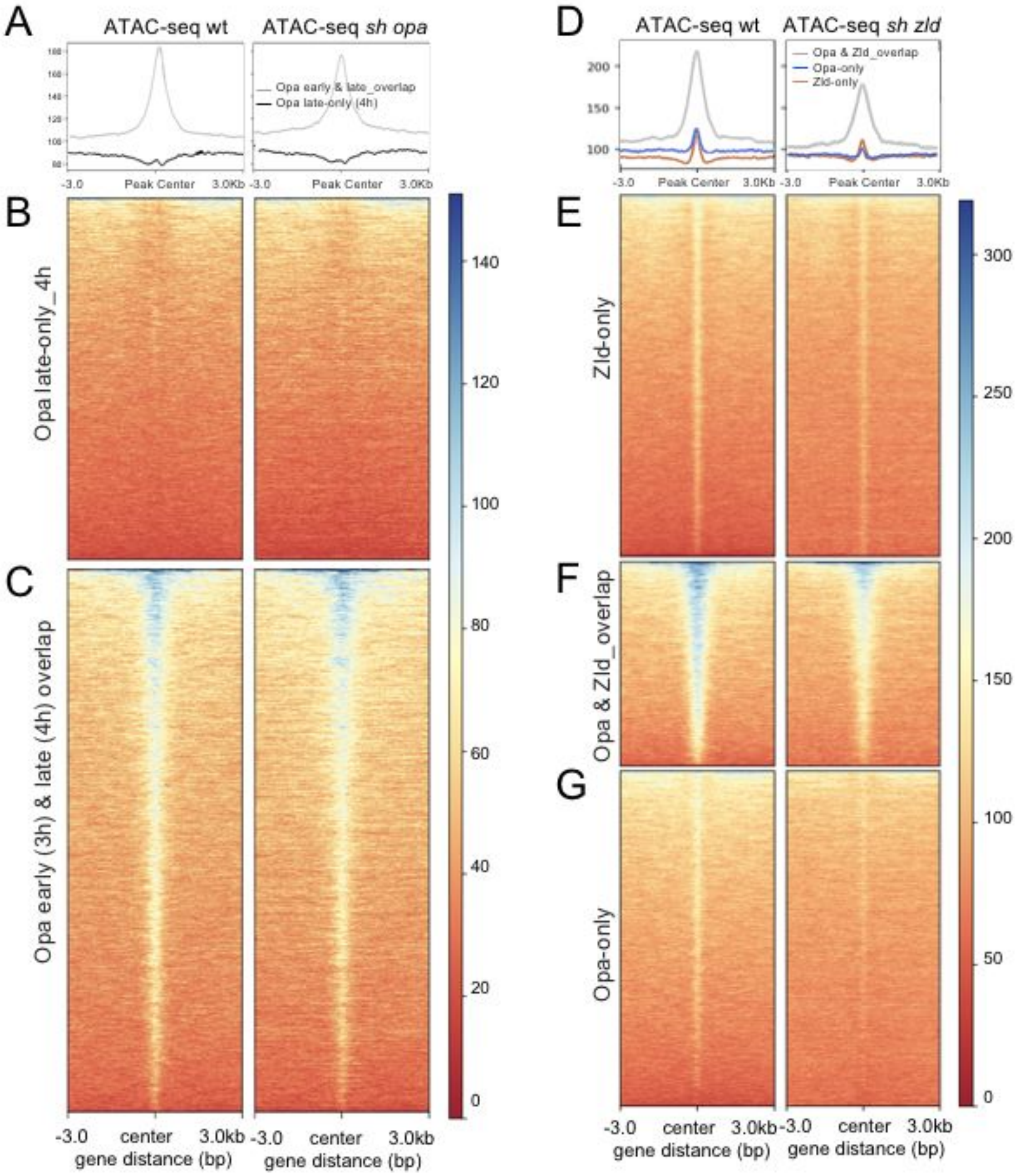


**Figure 5 - supplemental figure 1. ATAC-seq data for late-nc14 does not detect change in chromatin accessibility at sites of late Opa (4h) ChIP-seq regions but does confirm role for Zld at sites of early Opa (3h) ChIP-seq regions.**

(A-C) Aggregated signals and heatmaps of normalized ATAC-seq data from wt and *sh opa* mutant embryos for ChIP-seq Opa (4hr) Opa-only regions (top) compared to peaks in common between

Opa_early and Opa_late peaks called from ChIP-seq Opa-only (3hr) versus ChIP-seq Opa-only (4h) regions. Each line of the heatmap is a genomic region. The accessibility is summarized with a color code from red (no accessibility) to blue (maximum accessibility).

(D-G) Aggregated signals and heatmaps of late nc14 normalized ATAC-seq data from wt and *sh zld* mutant embryos for Zld-only (E; red trace, D), Zld & Opa overlap (F; gray trace, D) and Opa-only (G; blue trace, D). Another plot, on top of the heatmap, shows the mean signal at the genomic regions, which were centered to peaks of accessibility signals (D).

**Movie 1. Visualization of *sogD_Δopa* transcriptional activities in early embryo from nc12 to gastrulation u.**
